# Emergent behavioral organization in heterogeneous groups of a social insect

**DOI:** 10.1101/2020.03.05.963207

**Authors:** Yuko Ulrich, Mari Kawakatsu, Christopher K. Tokita, Jonathan Saragosti, Vikram Chandra, Corina E. Tarnita, Daniel J. C. Kronauer

**Affiliations:** Laboratory of Social Evolution and Behavior, The Rockefeller University, New York, NY 10065, USA; Department of Ecology and Evolution, University of Lausanne, 1015 Lausanne, Switzerland; Program in Applied and Computational Mathematics, Princeton University, Princeton, NJ 08544, USA; Department of Ecology and Evolutionary Biology, Princeton University, Princeton, NJ 08544, USA

**Author notes:** Corresponding author; (C.E.T.). Y.U. and M.K. contributed equally to this work. C.E.T. and D.J.C.K. contributed equally to this work. **Author Contributions** YU and DJCK conceived the study. MK, CKT, and CET developed the theoretical approach. YU and DJCK designed the experiments. YU, JS, and VC performed the experiments. YU analyzed the experiments. MK and CKT performed the simulations, and MK, CKT, and CET analyzed the simulation results. MK performed analytical calculations with input from CET. YU, MK, CKT, CET, and DJCK drafted the paper, and all authors provided comments.

**Keywords:** collective behavior, division of labor, self-organization, response threshold model, clonal raider ant

## Abstract

The composition of social groups has profound effects on their function, from collective decision-making to foraging efficiency. But few social systems afford sufficient control over group composition to precisely quantify its effects on individual and collective behavior. Here we combine experimental and theoretical approaches to study the effect of group composition on individual behavior and division of labor (DOL) in a social insect. Experimentally, we use automated behavioral tracking to monitor 120 colonies of the clonal raider ant, *Ooceraea biroi,* with controlled variation in three key correlates of social insect behavior: genotype, age, and morphology. We find that each of these sources of heterogeneity generates a distinct pattern of behavioral organization, including the amplification or dampening of inherent behavioral differences in colonies with mixed types. Theoretically, we use a well-studied model of DOL to explore potential mechanisms underlying the experimental findings. We find that the simplest implementation of this model, which assumes that heterogeneous individuals differ only in response thresholds, could only partially recapitulate the empirically observed patterns of behavior. However, the full spectrum of observed phenomena was recapitulated by extending the model to incorporate two factors that are biologically meaningful but theoretically rarely considered: variation among workers in task performance efficiency and among larvae in task demand. Our results thus show that different sources of heterogeneity within social groups can generate different, sometimes non-intuitive, behavioral effects, but that relatively simple models can capture these dynamics and thereby begin to elucidate the basic organizational principles of DOL in social insects.

**Significance Statement:** When individuals interact in an aggregate, many factors that are not known *a priori* affect group dynamics. A social group will therefore show emergent properties that cannot easily be predicted from how its members behave in isolation. This problem is exacerbated in mixed groups, where different individuals have different behavioral tendencies. Here we describe different facets of collective behavioral organization in mixed groups of the clonal raider ant, and show that a simple theoretical model can capture even non-intuitive aspects of the behavioral data. These results begin to reveal the principles underlying emergent behavioral organization in social insects. Importantly, our insights might apply to complex biological systems more generally and be used to help engineer collective behavior in artificial systems.

## Introduction

The study of collective behavior and self-organization is an active area of research across a diversity of fields, from animal movement (1) to robotics (2), from tissue engineering (3) to public health (4), and from voting (5) to conservation (6). The colonies of social insects in particular are striking examples of highly integrated, complex biological systems that can self-regulate without centralized control (7). Consequently, social insects have emerged as powerful systems to study collective behavior and social dynamics, both experimentally and theoretically (8–12). However, few experimental studies have comprehensively measured the influence of group composition—e.g., in age, genotype, or morphology—on collective organization, because the inherent complexity of many social insect colonies renders their composition intractable. This has limited our understanding of how colony composition affects both individual behavior and emergent group-level organization, and constitutes a major hurdle towards a general and comprehensive systems-level description of social insect colonies.

An emergent colony-level trait that has long been thought to depend on colony composition is division of labor (DOL). DOL describes the non-random variation in task performance among members of a social group (13), and is characterized both by between-individual variation and by individual specialization in task performance. Specifically, DOL has been hypothesized to increase with workforce heterogeneity, based on the observation that individual traits often correlate with individual task allocation (14). For example, workers of different age (15–18), genotype (e.g., patrilines (19, 20) or matrilines (21)), or morphology (e.g., size (19, 22–24)) can vary in their propensity to engage in tasks such as foraging, nursing, nest construction, or grooming of nestmates.

Experimentally testing this hypothesis in a systematic manner has proved challenging, even as theory has confirmed that workforce heterogeneity can indeed lead to the emergence of DOL (13, 25). One successful theoretical approach relies on the fixed response threshold model (FTM) of task allocation (26, 27). This model assumes that each task has an associated stimulus that signals the colony demand for that task. The magnitude of a given stimulus decreases with the number, efficiency, and/or average time investment of workers performing the corresponding task. Individuals respond to demands based on internal thresholds that reflect their sensitivity to the stimulus and govern their likelihood of performing a task given its stimulus level: the higher the stimulus level for a task relative to an individual’s threshold, the more likely it is to begin performing the task (see Materials and Methods for a detailed description). Thus, fixed thresholds provide a simple mechanism by which individuals dynamically allocate efforts to meet colony demands.

Previous work on the FTM has focused on differences in individual response thresholds as the primary driver of DOL (26, 28–32). In this simple formulation of the FTM, the heterogeneity in behavior is captured via heterogeneity in individual response thresholds drawn from a normal distribution with mean and variance that can be specific to the task and/or the type of individual. Yet, ants can also vary in other traits, for example in the efficiency with which they perform tasks (33–35) or in the average time spent performing a given task (36). Task demand can be similarly variable: for example, foraging activity levels of workers increase with the number of larvae that they have to tend to (37), and larvae of different genotypes develop into adults with different morphologies when cared for by the same workers (38). Thus, the level of demand emanating from the larvae could depend on their number and genotype. Despite this empirical evidence, few theoretical studies of DOL have explored the significance of inter-individual variation in traits other than response thresholds (14, 39).

Here we combine experimental and theoretical approaches to study the effect of group composition on both individual behavior and colony-level DOL. We use the FTM as a natural starting point, but systematically investigate a suite of parameters that might be associated with different individual traits of interest. To overcome the practical challenges associated with studying complex social systems empirically, we capitalize on the advantages of the clonal raider ant (*Ooceraea biroi*). The unique biology of this species affords unparalleled control over the main aspects of colony composition that are thought to affect individual- and group-level behavior in social insects: genotype, age, and morphology. Specifically, colonies of clonal raider ants are queenless and exclusively composed of workers that reproduce asexually and synchronously, so that all adults within a colony are genetically almost identical and emerge in discrete age cohorts. Furthermore, individuals show variation in ovariole number that is associated with body size and other morphological features (40), making it possible to approximately sort individuals into ‘regular workers’ (2-3 ovarioles) and ‘intercastes’ (4-6 ovarioles) based on their size (38). Conveniently, workers of different clonal genotypes, age cohorts, and morphologies can be mixed to create functional chimeric colonies (38). Taking advantage of these features, we quantify individual and collective behavior of *O. biroi* in response to precise, independent manipulations of colony composition along three independent axes, in a single system, and under standardized conditions.

## Results

### Baseline theoretical predictions of the ‘simple’ FTM with threshold heterogeneity

To establish baseline predictions in colonies with two types of ants (e.g., of different genotype, age, or morphology), we simulated experimental colonies using the simplest and most commonly-employed formulation of the FTM described above (see also Materials and Methods). Simulated colonies were either pure or mixed with respect to ant type; pure colonies consisted solely of one type of ant or the other, whereas mixed colonies had the two types in equal proportions. The ‘simple’ FTM assumes that the types only differ in mean response threshold. The individual thresholds for each type of ant are drawn from a normal distribution with the corresponding type-specific mean. All other model parameters—task performance efficiency, demand rate, threshold variance—are constant across types. Thus, the only source of heterogeneity in pure colonies was the distribution of individual response thresholds, while in mixed colonies that heterogeneity was compounded by differences in the means of the type-specific distributions. The assumption that some threshold heterogeneity exists even in pure colonies rests on the experimental observation that pure colonies exhibit DOL, yet in the absence of any type of heterogeneity, the FTM cannot produce DOL (32).

In pure colonies, there is a single normal distribution of individual thresholds for a given task. Because individuals from the lower end of the distribution are more sensitive to the stimulus for that task, they tended to perform that task more often than those from the higher end, resulting in DOL. In mixed colonies, there is a bimodal distribution of thresholds for each task, with the thresholds of the two types clustered around the different modes. This wider distribution of thresholds resulted in more pronounced DOL, i.e., both behavioral variation and specialization were greater in mixed colonies compared to pure colonies (Fig. 1a-b).

**Figure 1.**
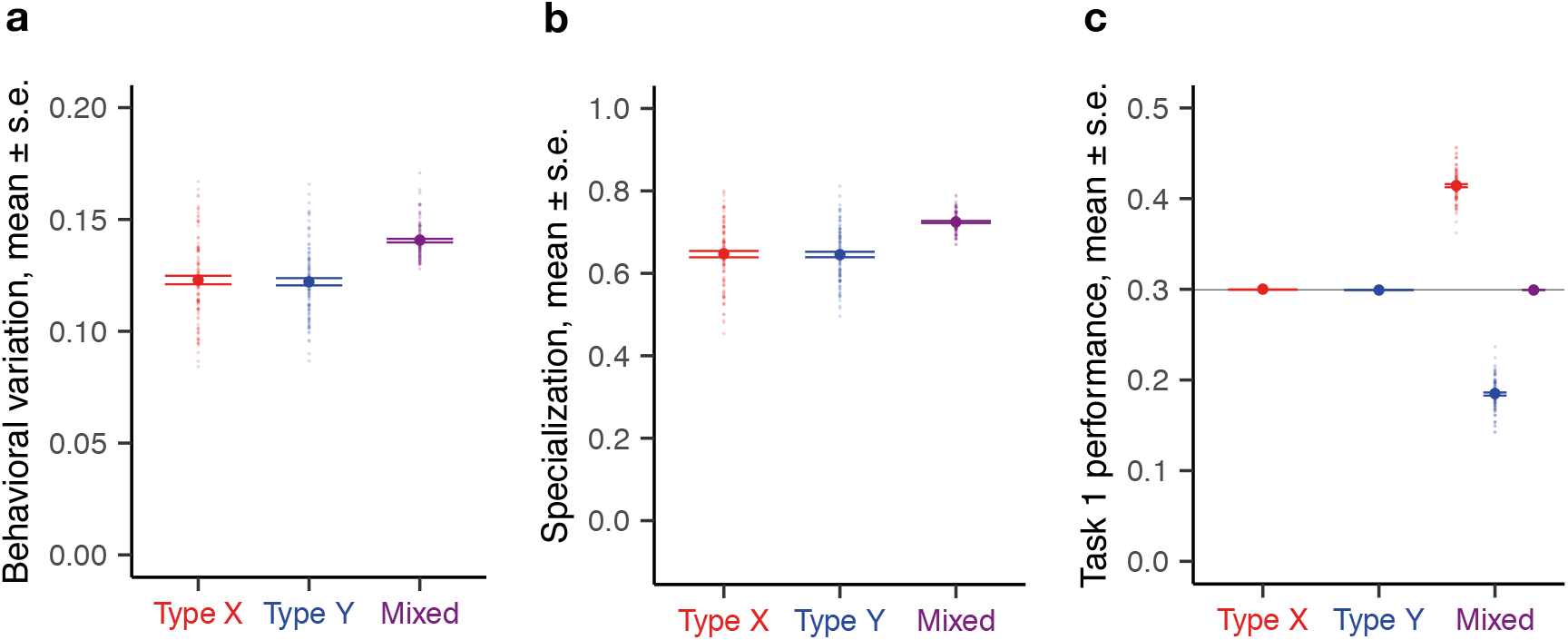
Theoretical predictions of the simple FTM with differences in mean task thresholds only. One hundred replicates were simulated for each colony composition. Each opaque circle represents an individual replicate colony (colony size 16); each solid circle represents the average value across all replicates for its corresponding colony (or sub-colony) composition. Panels show division of labor (behavioral variation (**a**), specialization (**b**)) and task performance frequency (**c**) as a function of colony composition. Type Y individuals have a higher mean threshold than type X individuals for both tasks (*μ*^*X*^ = 10, *μ*^*Y*^ = 20). All other parameters are identical for both types: *δ* = 0.6, *α* = 2, *σ* = 0.1, *η* = 7, *τ* = 0.2 (see Materials and Methods and Table S2 for parameter definitions).

However, all colonies, irrespective of their composition, had the same mean task performance (Fig. 1c). This is because, while colonies may differ in how they allocate workers to tasks (in this case, within mixed colonies, the two ant types differed in their mean task performance because the type with the lower average threshold for a given task took up that task more often than the other type), they must perform the same amount of work overall to satisfy a given demand. Thus, on average, colony members spent the same fraction of time performing each task across pure and mixed colonies.

In summary, the simple FTM predicted that (P1) regardless of composition, colonies would exhibit the same average task performance, but that (P2) mixed colonies would exhibit higher overall DOL and that (P3) the two types would behave differently from each other in mixed colonies, but not in their respective pure colonies (see Table 1 for a summary of predictions).

**Table 1.** Summary of theoretical results. Theoretical predictions of the simple FTM (top row) and extended FTM (bottom row) for pure and mixed colonies, as well as the pattern of behavioral change observed between them. Text in italic highlights key differences in model predictions. Colors indicate agreement (light green) or disagreement (light red) with experiments.

|  |  | Pure colonies |  | Mixed colonies |  | Behavioral change<br>from pure to mixed colonies |
| --- | --- | --- | --- | --- | --- | --- |
|  |  | Mean task performance | Division of labor (DOL) | Mean task performance | Division of labor (DOL) |  |
| Type of Fixed Threshold Model (FTM) | Simple FTM<br>(with variation in mean response threshold only) | *Identical across colonies of different ant types (P1)* | Exhibits DOL | *Identical to pure colonies (P1)* | Higher DOL than in pure colonies (P2) | Behavioral *amplification only* (P3) |
|  | Extended FTM<br>(with variation in other biologically relevant params) | *Different across colonies of different ant types (P4)* | Exhibits DOL | *Different from pure colonies (P4)* | Higher DOL than in pure colonies (P2) | Behavioral *contagion, amplification, or neither* (P5) |

= experiments **agree** with prediction = experiments **disagree** with prediction

### Effects of individual genotype, age, and morphology on individual behavior in experimental colonies

We experimentally tested these theoretical predictions in replicate experimental colonies that were either pure or mixed with respect to genetic, demographic, and morphological composition, manipulating each factor independently from the others (see Materials and Methods, Table S1). For example, demographically pure colonies contained either only young workers (1 month old) or only old workers (3 month old), and mixed colonies contained young and old workers in equal proportions; genotype and morphology were kept constant both within and between these colonies. Similarly, genetically pure colonies contained either only workers of genotype B or of genotype A (see (41) for genotype designations), and mixed colonies contained workers of the two genotypes in equal proportions; age and morphology were kept constant among these colonies. All colonies within an experiment had the same size. Colonies contained 8 or 16 workers—fully functional group sizes in the clonal raider ant—and the same number of age-matched larvae hosted in a Petri dish with a plaster floor (see Materials and Methods). The experiment on genetic effects was performed twice, once with larvae of each genotype.

We used a high-throughput automated tracking system (32) to record and analyze the behavior of all individual ants in 120 experimental colonies. The propensity of each ant to perform extranidal tasks (e.g., foraging, waste disposal) as opposed to intranidal tasks (e.g., nursing) was computed as the two-dimensional root-mean-square deviation (r.m.s.d.) of its spatial coordinates (32) (Fig. 2a; see Materials and Methods). The mean r.m.s.d of a group of ants was used as a proxy for their mean performance of extranidal tasks. To quantify colony-level DOL, we calculated behavioral variation and specialization among colony members. Behavioral variation was computed as the standard deviation across r.m.s.d. values of all ants from the same colony. Specialization was computed as the mean correlation between individual r.m.s.d. ranks across consecutive days in the experiment (32).

**Figure 2.**
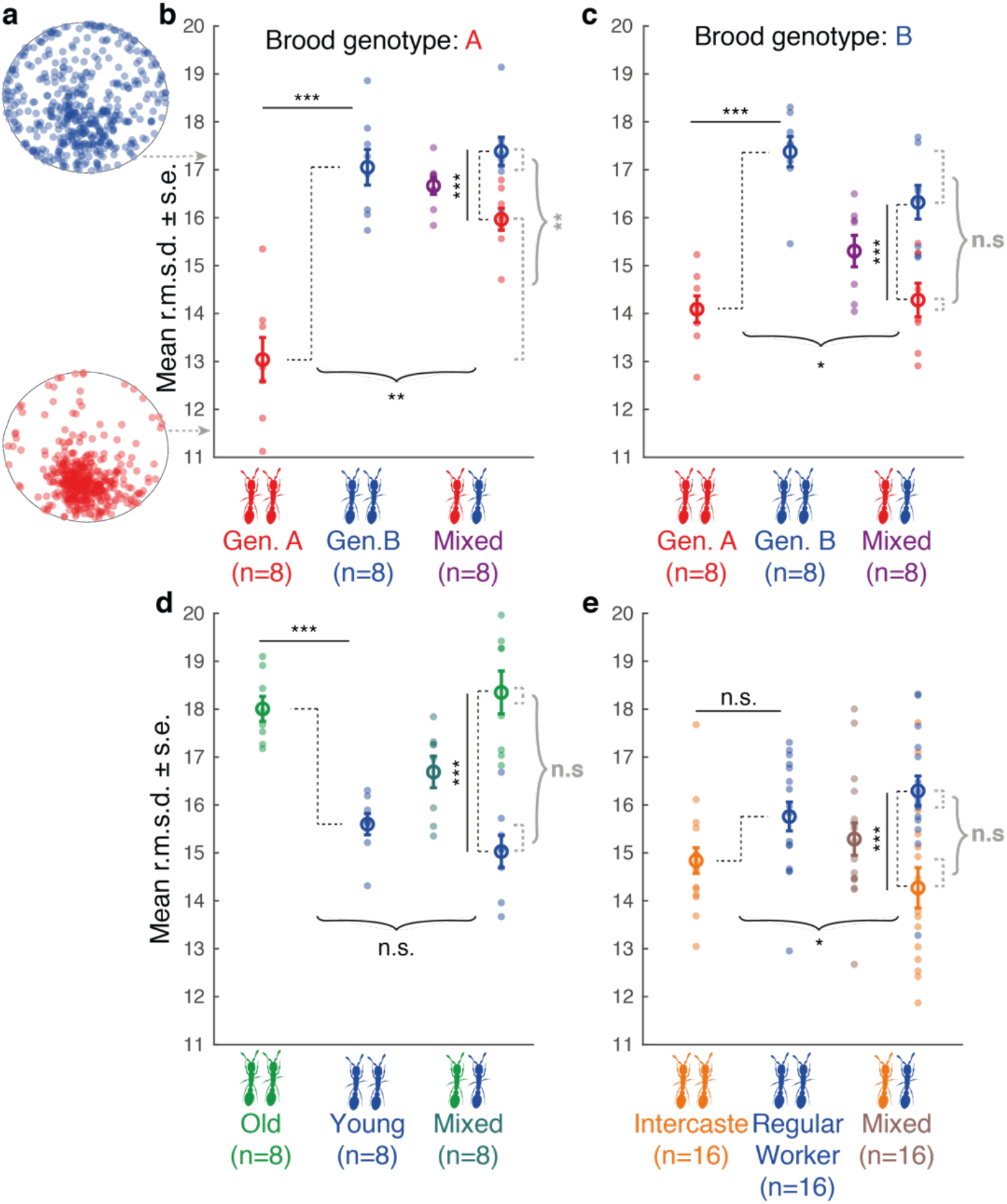
Mean r.m.s.d. (a proxy for mean extranidal activity) as a function of colony composition. Opaque circles represent mean behavior of individuals in replicate colonies (or sub-colonies). Open circles represent average values across replicate colonies (or sub-colonies). For mixed colonies, data are shown both as type-specific mean behavior (in type-specific colors) and colony-level mean behavior (in ‘average’ color). Identical colors across panels indicate ants of the same genotype, age, and morphological types. Sample sizes indicate the number of replicate colonies. Straight solid black brackets represent the effects of individual traits on behavior (X_p_ vs. Y_p_ and X_m_ vs. Y_m_). Black dotted brackets represent the behavioral differences between types in pure (Y_p_−X_p_) and mixed (Y_m_−X_m_) colonies. Black curly brackets represent the effect of mixing on inter-type behavioral differences (Y_p_−X_p_ vs. Y_m_−X_m_). Grey curly brackets represent the asymmetry of the effect of mixing between types (|X_p_−X_m_| vs. |Y_p_−Y_m_|). **a**: Spatial distribution of two ants with high (blue; genotype B) and low (red; genotype A) r.m.s.d. from the same colony. Arrows point to the corresponding r.m.s.d. values. **b:** Behavior as a function of colony genetic composition in colonies with A brood. Colony size 16. GLMM post hoc Tukey tests (B_p_ vs. A_p_: z = 7.75, p = 3.64*10^−14^; B_m_ vs. A_m_ : z = 4.61, p = 8.06*10-06) **c**: Behavior as a function of colony genetic composition in colonies with B brood. Colony size 16. (B_p_ vs. A_p_: z = 7.45, p = 2.80*10^−13^; B_m_ vs. A_m_ : z = 7.68, p = 6.57*10^−14^) **d:** Behavior as a function of colony demographic composition. Colony size 16 (Young_p_ vs. Old_p_: z = −6.05, p = 4.39*10^−09^; Young_m_ vs. Old_m_ : z = −13.31, p < 2*10^−16^). **e:** Behavior as a function of colony morphological composition. Colony size 8. (Regular Worker_p_ vs. Intercaste_p_: z = 2.14, p = 0.10, Regular Worker_m_ vs. Intercaste_m_ : z = 8.95, p < 2*10-16). n.s.: non-significant, *: p < 0.05, **: p < 0.01, ***: p < 0.001.

We found that workers of genotype B spent more time away from the nest (i.e., had higher mean r.m.s.d.) than workers of genotype A, both across pure colonies and within mixed colonies (Fig. 2b-c), suggesting a genetic basis for the propensity to perform extranidal tasks (19–21). Old workers spent more time away from the nest than young workers irrespective of colony demographic composition (Fig. 2d). Thus, *O. biroi* displays the classic form of age polyethism typical of social insects (15–18, 42), whereby older individuals allocate more time to extranidal tasks, and younger individuals spend more time at the nest. Finally, regular workers spent less time at the nest than intercastes in mixed colonies, but not across pure colonies (Fig. 2e). Because the larger body size and higher reproductive potential of intercastes correspond to a more queen-like phenotype, these behavioral differences support empirical data from other systems— including other queenless (43) and clonal (44) ant species—where reproductive potential often negatively correlates with foraging activity. Thus, consistent with existing knowledge, our experiments revealed robust differences in behavior (here, the propensity to perform extranidal tasks) across ant genotypes, age cohorts, and morphological types (Fig. 2). Interestingly, however, our experiments showed that different ant types (genotypes and age cohorts, but not morphologies) can have different mean behaviors between the corresponding two types of pure colonies. This is inconsistent with theoretical prediction (P1) that colonies, irrespective of their composition, should have the same mean behavior (Table 1).

### Effects of genetic, demographic, and morphological mixing on DOL and individual behavior in experimental colonies

We found that, in general, mixed colonies had higher DOL—measured as behavioral variation (Fig. S1) and specialization (Fig. 3)—than pure colonies. Although not all pairwise comparisons were statistically significant, there was no case where pure colonies had significantly higher DOL than mixed colonies. Thus, each of the three forms of workforce heterogeneity tended to promote DOL, consistent with prediction (P2) (see Table 1).

**Figure 3.**
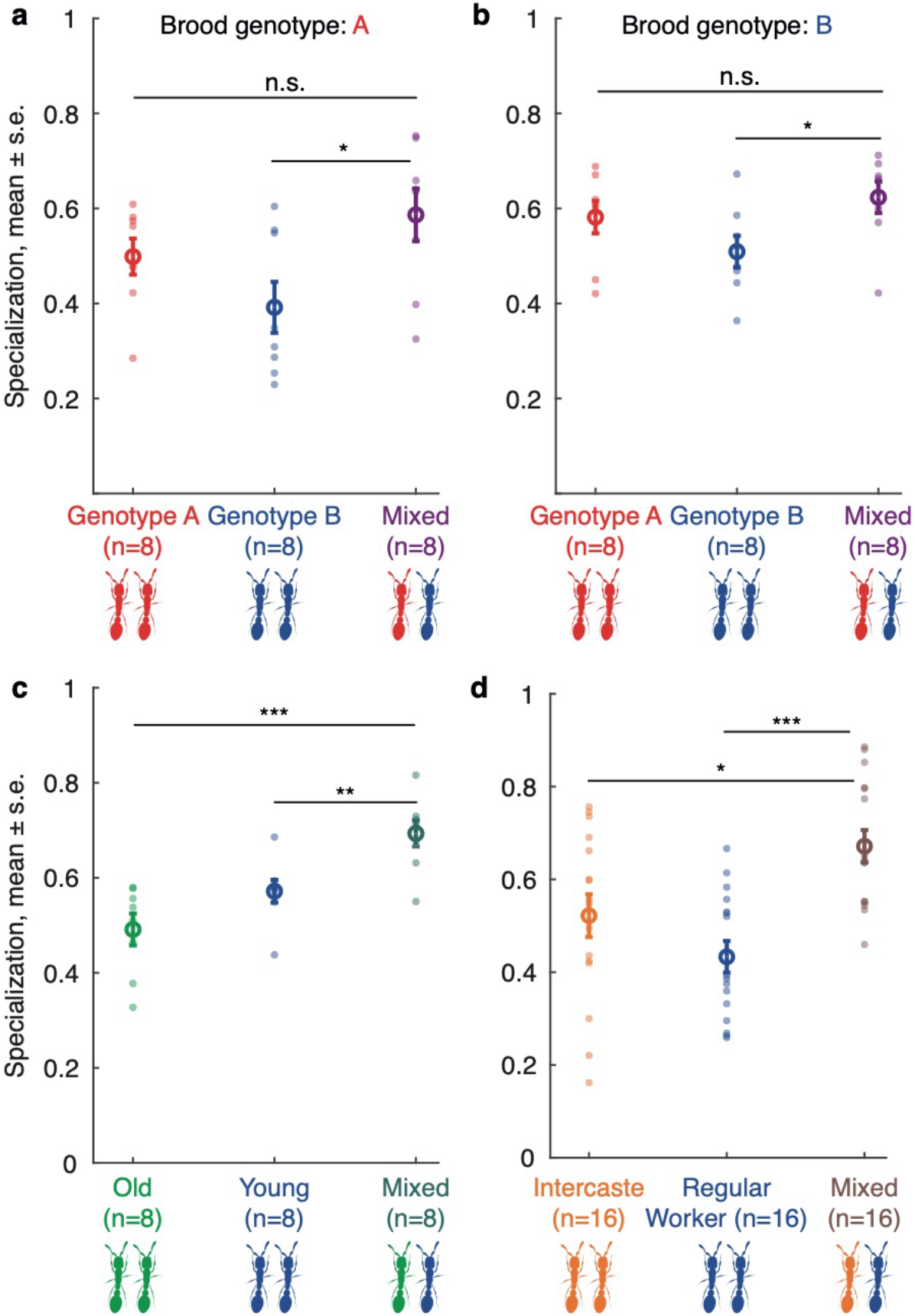
Specialization (day-to-day rank correlation in r.m.s.d.) as a function of colony composition. Opaque circles represent replicate colonies. Open circles represent average values across replicate colonies. Sample sizes indicate the number of replicate colonies. Identical colors across panels indicate ants of the same genotype, age, and morphological types. **a**: Specialization as a function of colony genetic composition in colonies with A brood. Colony size 16. (GLM post hoc Tukey tests; B_p_ vs. mixed: z = −2.78, p = 0.02; A_p_ vs. mixed: z = 1.25, p = 0.26) **b**: Specialization as a function of colony genetic composition in colonies with B brood. Colony size 16. (B_p_ vs. mixed: z = −2.41, p = 0.047; A_p_ vs. mixed: z = 0.88, p = 0.38) **c**: Specialization as a function of colony demographic composition. Colony size 16 (Young_p_ vs. mixed: z = 3.01, p = 0.005; Old_p_ vs. mixed: z = 5.01, p = 1.63*10^−06^) **d**: Specialization as a function of colony morphological composition. Colony size 8 (Regular Worker_p_ vs. mixed: z = −4.35, p = 4.05*10^−05^, Intercaste_p_ vs. mixed: z = 2.73, p = 0.013). n.s.: non-significant, *: p < 0.05, **: p < 0.01, ***: p < 0.001.

We next assessed the outcome of mixing individuals with different behavioral tendencies on individual behavior. Consider two types of individuals, X and Y. Let X_k_ and Y_k_ be the mean behavior of types X and Y, respectively, in pure (k = p) or mixed (k = m) colonies. We assume that Y_p_ > X_p_ and Y_m_ > X_m_, to reflect our observation that the type with higher r.m.s.d. in pure colonies always also had higher r.m.s.d in mixed colonies. Given this assumption, mixing could, in principle, have one of three possible outcomes on individual behavior:

1. No effect of mixing on individual behavior: the mean behavioral difference between types across pure colonies is the same as the mean behavioral difference between types within mixed colonies, so that Y_p_ − X_p_ = Y_m_ − X_m_.
2. Behavioral ‘contagion’: individuals of different types become behaviorally more similar on average to each other when mixed, so that Y_p_ − X_p_ > Y_m_ − X_m_; and
3. Behavioral ‘amplification’: individuals of different types become behaviorally more different on average from each other when mixed, so that Y_p_ − X_p_ < Y_m_ − X_m_.

The simple FTM predicted that the two different types will differ in mean behavior when mixed, but not when in pure colonies (P3) (Fig. 1c), i.e., that behavioral amplification should always be observed. However, in contrast to this theoretical prediction, all three outcomes were observed experimentally: genetic mixing resulted in behavioral contagion (Fig. 2b-c; Student’s *t*-test: *t* = 3.86, *p* = 0.002 in colonies with A brood, *t* = 2.62, *p* = 0.02 in colonies with B brood); demographic mixing had no effect on individual behavior (Fig. 2d; *t* = −1.50, *p* = 0.16); and morphological mixing resulted in behavioral amplification (Fig. 2e; *t* = −2.44, *p* = 0.02).

We further investigated whether mixing had an asymmetric effect on the two ant types, i.e., whether it affected one type more than the other, so that the magnitude of change in type-specific behavior between pure and mixed colonies was different across the two ant types (i.e. |X_m_ − X_p_| ≠ |Y_m_ − Y_p_|). Testing this hypothesis, we found evidence for asymmetric behavioral contagion in genetically mixed colonies with A brood (Fig. 2b), where mixing affected the behavior of A workers (by increasing their extranidal activity) more than it affected the behavior of B workers (*t*-test |A_m_ − A_p_| vs. |B_m_ − B_p_|: *t* = 3.86, *p* = 0.0024). All other scenarios studied displayed symmetric effects of mixing on individual behavior (Fig. 2c: |A_m_ − A_p_| vs. |B_m_ − B_p_|: *t* = −0.94, *p* = 0.37; Fig. 2d, |Young_m_ − Young_p_| vs. |Old_m_ − Old_p_|: *t* = −1.02, *p* = 0.33, Fig. 2e, |Regular Worker_m_ − Regular Worker_p_| vs. |Intercaste_m_ − Intercaste_p_|: *t* = 0.68, *p* = 0.50).

Thus, both the direction and the magnitude of change in individual behavior between pure and mixed colonies depended on the specific source of workforce heterogeneity.

### Theoretical predictions of the extended FTM

The predictions of the simple FTM only partially captured the patterns observed in the experimental colonies (Table 1). Thus, differences in mean threshold alone were insufficient to explain the observed data, suggesting the need to consider other biologically realistic sources of heterogeneity in the model.

Much like assuming that types differ solely in their threshold means, assuming that types differ only in threshold variance or duration of task performance failed to capture the experimentally observed difference in mean behavior between pure colonies (Fig. S2a-b). However, between-type differences in task performance efficiency alone did reproduce this difference. In fact, if the demand was the same for both tasks, differences in task efficiency were necessary for such a pattern to emerge (SI Appendix).

When types differed only in task performance efficiency, we further found behavioral contagion in mixed colonies, i.e., the types behaved more similarly to each other when mixed. Critically, the asymmetry of this contagion depended on the magnitude of the task demand. If the task demand was not too high, so that both types could keep up with the demand in their pure colonies, then the contagion was always downward (Fig. 4a; analytical results in SI Appendix), i.e., the mixed colony, on average, behaved more like the more efficient type. If, on the other hand, the task demand was so high that the less efficient type could not keep up with task demand in its pure colony, then the contagion could, for certain parameter combinations, be upward (Fig. 4b), i.e., the mixed colony behaved on average more like the less efficient type. Hence, if in addition to differences in task efficiency we also assumed between-type differences in task demand (to reflect possible differences in the intensity of task demand stemming from larvae of different genotypes), we were able to qualitatively recapitulate the asymmetric behavioral contagion observed in genetically mixed colonies. Holding all else fixed, differences in task efficiency guaranteed behavioral contagion; the magnitude of task demand modulated the asymmetry of this contagion, i.e., whether mixed colonies on average behaved more like the more or less efficient type.

**Figure 4.**
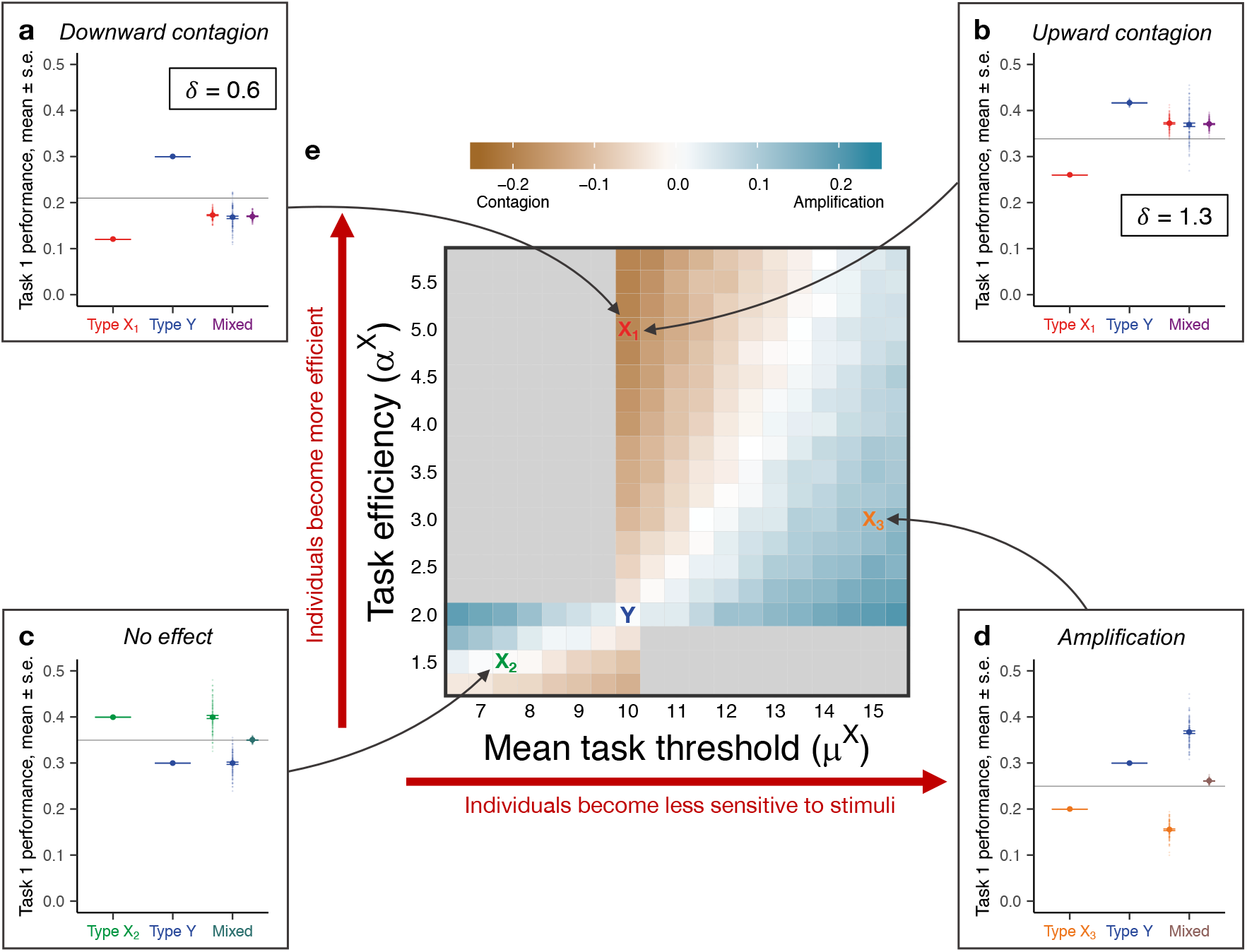
Theoretical predictions of the FTM on task performance and their robustness. **a-d:** Task performance frequency as a function of colony composition. One hundred replicates were simulated for each colony composition. Each opaque circle represents a replicate colony (colony size 16); each solid circle represents the average value across all replicates for its corresponding colony (or sub-colony) composition. Horizontal gray lines represent the average value of the pure colonies (first two columns) in their respective panels. Identical colors across panels indicate ants of the same types; in particular, the parameters for type Y ants are fixed across panels **a-d** (*μ*^*Y*^ = 10, *α*^*Y*^ = 2). **a-b**: Differences in task efficiency (α) between types and demand rate (*δ*) across colonies capture asymmetric behavioral contagion, downward (**a**) and upward (**b**). Larvae are more demanding in **b** (*δ* = 1.3) than in **a** (*δ* = 0.6). For a given *δ*, type X_1_ is more efficient than type Y (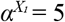, *α*^*Y*^ = 2). Type Y can keep up with demand when the larvae are less demanding (**a**) but not when they are more demanding (**b**); type X_1_ can keep up with the demand in both cases. Parameters: *σ* = 0.1, *μ* = 10, *η* = 7, *τ* = 0.2. **c-d**: Between-type differences in task efficiency (α) and mean threshold (*μ*) capture both a lack of effects from mixing (**c**) and behavioral amplification (**d**). In **c**, type X_2_ is less efficient than type Y (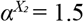, *α*^*Y*^ = 2) and has a lower threshold for both tasks (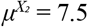, *μ*^*Y*^ = 10). In **d**, type X_3_ is more efficient than type Y (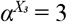, *α*^*Y*^ = 2) and has a higher threshold for both tasks (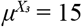, *μ*^*Y*^ = 10). Parameters: *σ* = 0.1, *η* = 7, *δ* = 0.6, *τ* = 0.2. **e:** Change in between-type relative task task performance between mixed and pure colonies (measured as (Y_m_−X_m_) − (Y_p_−X_p_)) as a function of type X’s task efficiency, *α*^*X*^, and mean task threshold, *μ*^*X*^. The letters X_1_, X_2_, and X_3_ indicate the parameter settings for type X in **a-d**; the blue letter Y indicates the parameter settings for type Y, which are fixed for **a-d** and all grids in **e**. Shades of blue indicate behavioral amplification (Y_p_−X_p_ < Y_m_−X_m_), and shades of brown indicate behavioral contagion (Y_p_−X_p_ > Y_m_−X_m_); light gray indicates regions in which the behavior is undefined according to our definitions of the behavioral patterns, which exclude biologically unrealistic scenarios (see Results). Fifty replicates were simulated for each parameter combination. Parameters: *η* = 7, *σ* = 0.1, *τ* = 0.2.

Although the combination of between-type differences in efficiency and demand successfully recapitulated the observed behavioral contagion, it failed to capture the other observed effects of mixing on individual behavior, notably instances where mixing had no effect on behavior, or where it resulted in behavioral amplification. If we instead combined the between-type differences in task efficiency with between-type differences in mean threshold (to reflect possible between-type differences in the intrinsic propensity to perform tasks), we were able to qualitatively recapitulate both the effects of demographic mixing (no effect of mixing on type-specific behavior; Fig. 4c) and the effects of morphological mixing (behavioral amplification in mixed colonies; Figs. 4d, S3). Whether we recapitulated the former or the latter depended on the magnitude of the difference in mean thresholds: a larger difference caused the types to differentiate their behavior more strongly in mixed colonies, leading to the latter; a smaller difference dampened this effect, leading to the former.

In general, the model robustly produced a spectrum of patterns, from behavioral contagion to amplification, across a large parameter space (Fig. 4e). Thus, incorporating additional, biologically realistic sources of heterogeneity into the model led to predictions that qualitatively mirrored the range of empirically observed behavioral patterns, namely: (P4) pure colonies of different ant types can differ from each other and from the mixed colonies in mean behavior, and (P5) mixing two types of ants can lead to behavioral contagion, amplification, or neither (Table 1). Moreover, these extensions preserved prediction P2 (to the extent observed in the experimental data), that mixed colonies tend to have higher DOL than pure colonies (Figs. S4, S5).

### Theoretical predictions for mean task performance in non-1:1 mixes

Despite its simplicity, the fixed threshold framework demonstrated remarkable explanatory power in both pure and mixed colonies. Given this success, we used the extended FTM to further explore expected patterns of task allocation in colonies with different ratios of ant types. We focused on the four parameter combinations in Fig. 4 because our analysis showed that they collectively captured all of the patterns observed in the experiments. For each parameter combination, we investigated how the mean task performance of colonies changed as we varied the ratio of the two ant types.

Simulations predicted a striking range of patterns. For the parameter combination that produced no effect in the mixed colonies with equal proportions of the two ant types (‘1:1 mixes’), the model produced an approximately linear relationship between mean task performance and the ratio of ant types (Fig. 5a). In all other cases, the mean task performance depended nonlinearly on the ratio of the types. However, the shape of the nonlinear curve differed among the cases. In the cases corresponding to behavioral contagion in the 1:1 mixes, the relationship followed a convex decreasing function, so long as there were enough individuals of the more efficient type such that the colony could keep up with the demand (Fig. 5b; analytical results in SI Appendix); otherwise the colony performed the tasks at a fixed maximum capacity that depended only on the average task duration (Fig. 5c). In the case corresponding to behavioral amplification, the relationship followed a concave decreasing function (Fig. 5d). Hence, despite one type being more efficient than the other in all cases considered, replacing an individual of the former type with one of the latter type would lead to qualitatively different outcomes depending on the between-type differences in mean threshold.

**Figure 5.**
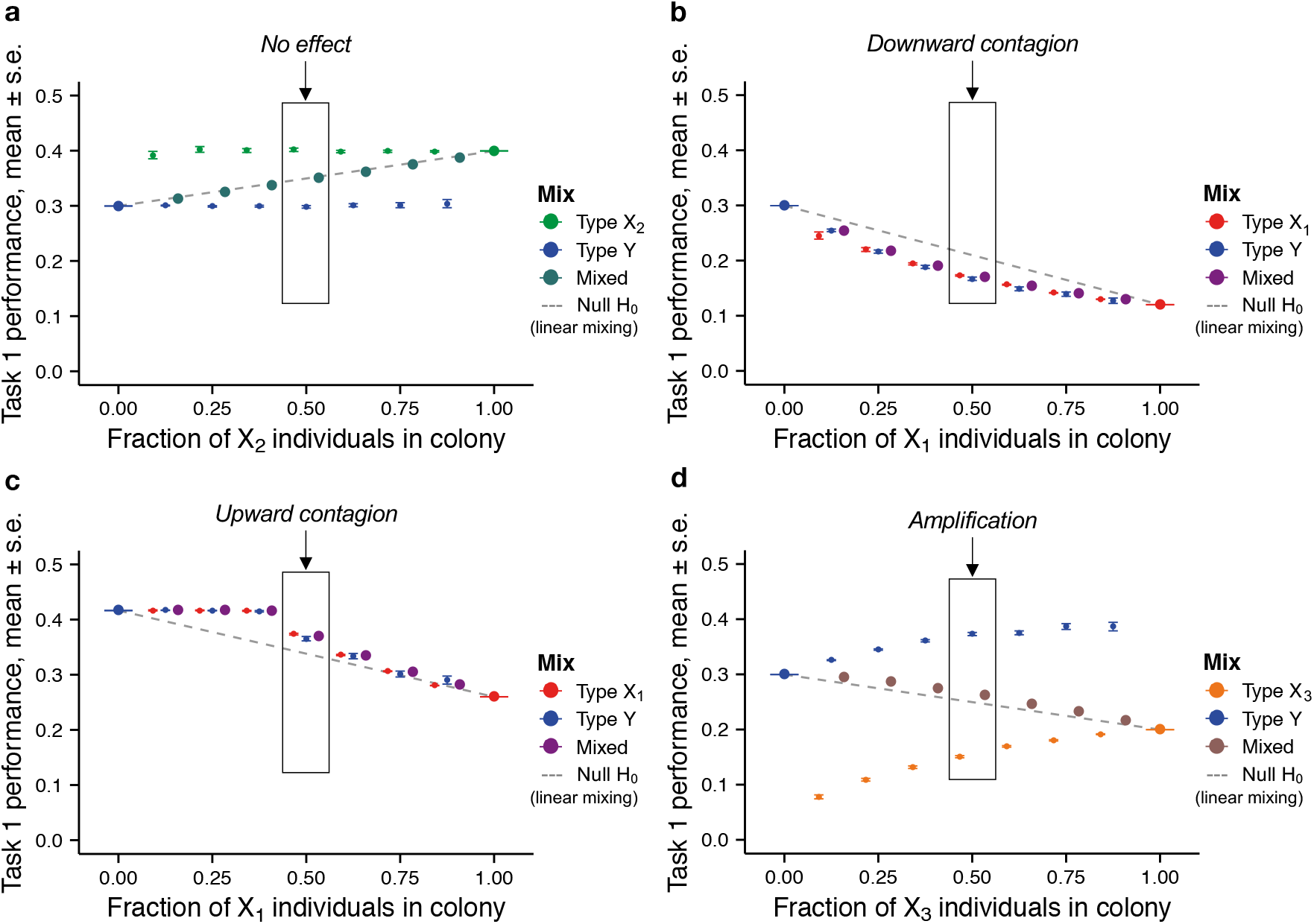
Predictions of the FTM for non-1:1 mixes. Colonies of size 16 with varying ratios of X and Y individuals were simulated under different conditions of threshold values, task-performance efficiency, and task demand. One hundred replicates were simulated for each colony composition. Each large circle represents the mean for that mix of X and Y individuals, while the neighboring smaller circles represent the means of X and Y individuals, respectively, within that mix. The dashed lines indicate the null hypothesis of linear behavioral effects of mixing types. The boxes highlight the behavioral patterns that characterize the 1:1-mixes, and their labels indicate correspondence with panels in Fig. 4 (**a** with Fig. 4c, **b** with Fig. 4a, **c** with Fig. 4b, and **d** with Fig. 4d). Parameters for each type (X_1_, X_2_, X_3_, Y) are identical to those of the corresponding type in Fig. 4. **a**: X_2_ individuals have a lower mean task threshold and are less efficient than Y individuals. **b**: X_1_ individuals are more efficient than Y individuals. **c**: X_1_ individuals are more efficient than Y individuals, but task demand is high. **d**: X_3_ individuals have a much higher mean task threshold than Y individuals and are more efficient.

Regardless of the case studied, the ratio of the types did not alter the qualitative effect of mixing on individual behavior (behavioral contagion, amplification, or no effect); for example, the case that led to behavioral amplification in 1:1 mixes predicted behavioral amplification for all non-1:1 mixes tested (Fig. 5d).

## Discussion

By manipulating social group composition along three different axes, we found that the effects of group heterogeneity on behavioral organization vary qualitatively depending on the specific factor under consideration. When ants of two different genotypes, ages, or morphologies were mixed, the inherent behavioral differences between each pair of types were dampened, unaffected, or amplified, respectively. The fact that various sources of heterogeneity that naturally exist in animal groups can have different, and possibly opposing, effects on collective organization underscores the importance of independently considering and controlling them. In nature, as in many experiments, all the factors studied here (larval and worker genotype, age, and morphology), as well as other effects (e.g., environmental conditions, resource availability) will play out simultaneously and in largely intractable ways. Being able to break this complexity down experimentally to study each effect separately and under standardized conditions is unprecedented and provides new insight into the basic organizing principles of behavior in social groups.

The experimental literature on DOL in social insects has historically attributed most inter-individual variation in behavior to variation in response thresholds. Our combined empirical and theoretical analyses, however, suggest that this is only part of the story. Indeed, we found that the simplest and most common implementation of the FTM, which assumes that individuals vary in response thresholds alone, only explained part of the empirically observed patterns of behavior. However, the full spectrum of observed phenomena could be qualitatively recapitulated by extending the model to incorporate heterogeneity in two additional factors: task performance efficiency and task demand. Both are empirically documented (33–35, 45, 46) but theoretically rarely considered.

Between-type differences in threshold and in task efficiency alone—two sources of heterogeneity with opposing effects on behavioral organization—were sufficient to recapitulate the core of our empirical results. Between-type differences in threshold led to behavioral amplification, making ant types behaviorally more different when mixed than when separated, as is known from previous theoretical work on the FTM (26). In contrast, between-type differences in task efficiency led to behavioral contagion, making ant types behaviorally more similar when mixed than when separated. Our theoretical analysis suggests that the relative strengths of these two sources of heterogeneity might vary with colony composition. In our experiments, varying colony *morphological* composition produced behavioral patterns that were theoretically recapitulated under a relatively strong effect of between-type differences in threshold and a relatively weak effect of differences in efficiency. In contrast, the behavioral patterns observed under varied *genetic* composition matched the theoretical predictions for the case in which differences in efficiency have a relatively stronger effect. Manipulating *demographic* composition corresponded to an intermediate scenario in which the two opposing forces seemed to balance each other out. While both threshold (47–49) and efficiency (33–35) are known to vary with various individual traits in social insects, their relative contributions to age-, genotype- and morphology-based behavioral variation remain poorly understood and deserve further investigation.

A third source of heterogeneity, task demand, was necessary to recapitulate the asymmetry in behavioral contagion (i.e., whether workers in mixed colonies behaved more like one or the other type of workers in pure colonies). Empirically, whether a colony composed of two ant genotypes behaved more like one genotype or the other depended on the genotype of the larvae reared. Coupled with the theoretical analysis, these results suggest that the differences in brood genotype could be a source of differences in task demand. This points to the brood as an important player in the regulation of task allocation, at least for tasks associated with brood care, such as foraging and nursing (50). That the brood can influence colony-level traits has been shown in several social insects where larvae solicit food from workers via chemical (51–53) or behavioral (54, 55) cues that affect worker physiology (52, 56) and behavior (e.g., foraging (57), feeding (58)). However, the effect of larvae on the allocation of tasks across individual workers remains elusive in many social insects due to the challenges associated with measuring individual behavior in groups and precisely controlling brood demand. By taking advantage of automated tracking and the unique biology of the clonal raider ant, our study overcomes these challenges and advances our understanding of larval factors that affect task allocation: we suggest that brood demand and its effects on task allocation depend not only on the presence and number of larvae (37, 59), but also on larval genotype. These results also provide insights into previous cross-fostering experiments that revealed that interactions between worker and brood genotypes have non-linear effects on brood development (into intercastes vs. regular workers) (38). Our work suggests that these effects might arise, at least in part, because different larval genotypes signal different levels of demand—and thereby differ in the magnitude of their effect on worker behavior—while different worker genotypes differ in their behavioral response to a given level of larval demand. For example, if different larval genotypes solicit food at different rates and different worker genotypes respond differently (e.g., via foraging thresholds or efficiency) to such differences in demand, the interaction between genetically-based larval demand and worker behavioral responses may result in differences in larval nutrition. Such differences may, in turn, lead to the previously reported shifts in larval development and, therefore, adult phenotype (38).

Overall, these findings demonstrate that, despite its simplicity, the FTM has remarkable versatility in recapitulating a broad range of experimental outcomes, while still operating under biologically plausible assumptions. It is important to note, however, that while the behaviors observed are robust and generic—i.e., the parameters chosen to illustrate the versatility of the FTM are representative of large regions of parameter space—little is known about what parameter values might actually correspond to the different experimental types. Nevertheless, even in the absence of such experimental measurements, the model provides a useful starting point to generate testable predictions for increasingly complex colony compositions in the clonal raider ant and possibly other social insects.

Our findings add to the growing literature on the role of individual heterogeneity in the collective behavior of complex biological (e.g., schools of fish, neurons in a brain, pathogen strains sharing a host, etc.) and artificial (e.g., heterogeneous robot swarms, synthetic microbial communities, etc.) systems. Much like colonies of the clonal raider ant, these systems exhibit patterns that can be interpreted as behavioral convergence (60–64), divergence (65), and non-linear effects of mixing on group-level phenotypes (66–68). In turn, these patterns affect important processes such as collective decision-making (5), the transmission and evolution of disease (69, 70), and the evolution of cooperative behavior (71, 72). While different variants of threshold-based models have been employed to study several of these systems (73–76), we still lack a unified theoretical framework to understand the consequences of individual differences on collective dynamics (77). Thus, a comparative approach to the study of the basic organizing principles of heterogeneous systems across scales constitutes an important next step towards understanding the behavior of complex biological systems.

## Materials and Methods

### Experimental design

Four experiments were performed to investigate the effect of genetic composition (2 experiments differing in the brood genotype used), demographic composition (1 experiment), and morphological composition (1 experiment). Each experiment comprised three treatments (2 with pure colonies, 1 with mixed colonies). All colonies within one experiment were monitored in parallel, but the different experiments were performed separately.

Experimental colonies were composed of workers of the desired age, genotype and morphology (Table S1), as well as larvae, housed in airtight Petri dishes 5 cm in diameter (corresponding to about 25 ant body-lengths) with a plaster of Paris floor. To control individual genotype, clonally related workers were sourced from the same stock colony. We used two commonly used genotypes, A and B (32, 38, 78). To control individual age, workers were sourced from a single age cohort from the same stock colony. Owing to the synchronized reproduction of *O. biroi*, all age-matched workers collected this way had eclosed within a day of each other. To control individual morphology, age-matched regular workers and intercastes from the same stock colony were screened based on body size (small or large) and the absence or presence of vestigial eyes, respectively. From the time they were collected (1–3 days after eclosion) until the start of experiments, workers of a given type were kept as a group. All workers were tagged with color marks on the thorax and gaster using oil-paint markers. Experimental colonies contained 16 (genetic composition and demographic composition experiments) or 8 (morphological composition experiment) workers and a matching number of age-matched larvae (4-5 days old). This 1:1 larvae-to-workers ratio corresponds to the estimated ratio found in a typical laboratory stock colony in the brood-care phase. We used 8 (genetic composition and demographic composition experiments) or 16 (morphological composition experiment) replicate colonies were set up for each group composition, for a total of 120 colonies.

The experiments took place in a climate room at 25 °C and 75% relative humidity under constant light (*O. biroi* is blind and its behavior is not affected by light). Every 3 days, we cleaned and watered the plaster, and added one prey item (live pupae of fire ant minor workers) per live larva at a random location within the Petri dish.

Behavioral data acquisition and analyses were performed as in (32). Software for image analysis is available at https://doi.org/10.5281/zenodo.1211644.

### Behavioral data analyses

*O. biroi* colonies switch between reproductive phases, in which all workers stay in the nest and lay eggs, and brood-care phases, in which workers nurse the larvae in the nest but also leave the nest to forage, explore, or dispose of waste. For each colony, behavioral analyses were restricted to the brood-care phase, which started at the beginning of the experiment and ended when all larvae had either reached the non-feeding pre-pupal stage or died.

The spatial distribution of each ant throughout the brood-care phase was quantified as the two-dimensional root-mean-square deviation:

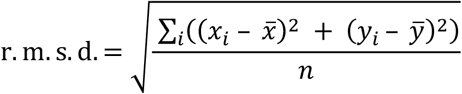

in which *x*_*i*_ and *y*_*i*_ are the coordinates of the focal ant in frame *i*, and *x̅* and *y̅* are the coordinates of the center of mass of the focal ant’s overall spatial distribution in the brood-care phase, and *n* is the number of frames in which the focal ant was detected. The r.m.s.d. is bounded between 0 and *r*, the radius of the Petri dish. Workers that spend a lot of time at the nest with the brood (e.g., nursing the larvae) and little time performing extranidal tasks (foraging or waste disposal) have low r.m.s.d. values, whereas workers that spend more time away from the brood have higher r.m.s.d. values.

For each colony, mean behavior was computed as the average of individual r.m.s.d. values, and behavioral variation was computed as the standard deviation of individual r.m.s.d. values. Both metrics were then averaged across replicate colonies for each treatment.

To quantify specialization, we use a metric appropriate for use on continuous behavioral data (r.m.s.d.). Specialization was defined for each colony as the Spearman correlation coefficient between individual r.m.s.d. ranks on consecutive days, averaged over the brood-care phase. Mean rank-correlation coefficients were then compared across treatments. For all behavioral analyses, ants were excluded from the dataset if they were detected in less than 30% of the frames acquired within the considered time frame (brood-care phase or day); for ants that died during the brood-care phase, the considered time frame was the portion of the brood-care phase preceding death.

### Statistical analyses

Statistical analyses were performed in R (79). Analyses were performed separately for each of the four experiments. As the experiments were performed at different times using different cohorts of ants, we cannot rule out “batch” effects and therefore avoid any statistical analyses comparing treatments across experiments.

#### Effects of individual attributes traits on behavior

The effects of colony composition (pure, mixed), individual attributes (genotypes A vs. B, Young vs. Old, or Regular worker vs. Intercaste), and their interaction, on individual behavior (individual r.m.s.d.) were investigated using linear mixed effects (LME, function *lmer* of package *lme4*) models with colony as a random factor. If a significant interaction between colony composition and individual attributes was detected, we used a second LME model with a four-level independent fixed variable combining colony composition and individual attributes (X_p_, Y_p_, X_m_ and Y_m_, where X_k_ and Y_k_ are the mean behavior of ant types X and Y, respectively, in pure (k=p) or mixed colonies (k=m)), followed by a Tukey posthoc test with Bonferroni-Holm correction (function *glht* of package *multcomp*) for the following planned comparisons: X_p_ vs. X_m_, Y_p_ vs. Y_m_, X_p_ vs. Y_p_, and X_m_ vs. Y_m_. When needed, response variables were transformed to satisfy model assumptions.

#### Effects of genetic, demographic, and morphological mixing on DOL

The effects of the treatment (a 3-level variable: X, Y, and mixed) on division of labor (behavioral variation, specialization) were investigated using generalized linear models (GLM), followed by Tukey posthoc tests with Bonferroni-Holm correction for all three pairwise comparisons.

#### Effects of genetic, demographic, and morphological mixing on individual behavior

To assess whether type-specific behavior was affected by colony composition, we compared the difference in mean behavior (mean r.m.s.d.) between types across pure colonies to the difference in mean behavior between the same types within mixed colonies (i.e., Y_p_ − X_p_ vs. Y_m_ − X_m_, where Y_p_ > X_p_ and Y_m_ > X_m_), using unpaired t-tests, after verifying assumptions of normality. We further tested whether the amplitude of the effect differed across types by comparing the magnitude of change in type-specific behavior between pure and mixed colonies across the two ant types (i.e. |X_m_ − X_p_| ≠ |Y_m_ − Y_p_|) with unpaired t-tests, after verifying assumptions of normality.

### Theoretical model

The fixed threshold model (FTM) considers a colony of *n* individuals, *N*_*X*_ of which are of type X and *N*_*Y*_ are of type Y (*N*_*X*_ + *N*_*Y*_ = *n*). Types X and Y represent any pair of the experimentally manipulated sub-colony compositions (i.e., genotypes A and B, Young and Old, or Regular Workers and Intercastes). Without loss of generality, we assume that individuals 1, …, *N*_*X*_ are of type X and individuals *N*_*X*_ + 1, …, *n* are of type Y. The colony must perform *m* tasks; for consistency with the experimental approach, we assume that there are two tasks (*m* = 2). At a given time step, an individual can be either performing one of the *m* tasks (active) or not performing any (inactive). The *task state* of individual *i* at time *t* is given by the binary variable *x*_*ij,t*_: if individual *i* is active and performing task *j* at time t, then *x*_*ij,t*_ = 1 and *x*_*ij’,t*_ = 0 for all *j’* ≠ *j*; if individual *i* is inactive and in its rest state, then *x*_*ij,t*_ = 0 for all *j*. The task state of the colony at time *t* is then given by the *n*-by-*m* binary matrix *Q*_*t*_ = [*x*_*ij,t*_].

The FTM further assumes that each task *j* has an associated *stimulus s*_*j,t*_. This stimulus signals the group-level demand for task *j* and changes depending on both the rate at which the demand increases (e.g., the demand for foraging increases due to increased hunger in the colony) and the number of individuals performing the task (e.g., the demand for foraging decreases as workers go out and find food). Mathematically, the change in stimulus *s*_*j,t*_ is governed by Eq. (1):

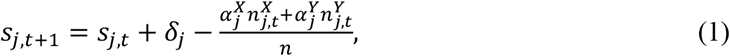

where *δ*_*j*_ is the task-specific demand rate, taken to be constant over time; *α*_*j*_^*X*^ and *α*_*j*_^*Y*^ are the task-specific performance efficiency of type X and Y individuals, respectively; and 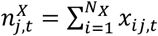 and 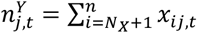 are the numbers of type X and Y individuals performing task *j* at time *t*, respectively.

Each individual *i* is assumed to have an internal *threshold* for task *j*, *θ*_*ij*_, drawn at time *t* = 0 from a normal distribution with mean *μ*_*j*_ and normalized standard deviation *σ*_*j*_ (i.e., expressed as a fraction of the corresponding mean *μ*_*j*_). Although thresholds may change over the individuals’ lifetime, they are assumed to be fixed over the timescale of the experiments and, consequently, over the simulation runs. We refer to *μ*_*j*_ as the *mean task threshold* and to *σ*_*j*_ as the *threshold variance* for task *j*; each can be group- and/or task-specific (i.e., 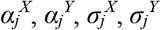).

At each time step, inactive individuals assess the *m* task stimuli in a random sequence until they either begin performing a task or have encountered all stimuli without landing on a task. For each encountered stimulus, individual *i* evaluates whether to perform the task by comparing the stimulus level to its internal threshold. Specifically, given stimulus *s*_*j,t*_ and internal threshold *θ*_*ij*_, individual *i* commits to performing task *j* with probability

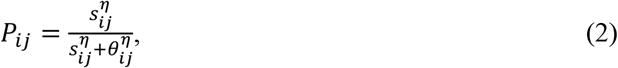

where parameter *η* governs the steepness of this response threshold function. The larger the value of *η*, the more deterministic the behavior; in the limit *η* → ∞, the response function becomes a step function (*H*[*s*_*j,t*_ − *θ*_*ij*_] = 0 if *θ*_*ij*_ > *s*_*j,t*_, 1 if *θ*_*ij*_ < *s*_*j,t*_) where *H* is the Heaviside function). Active individuals spontaneously quit their task with a constant quit probability *τ*. Active individuals can neither evaluate stimuli nor switch tasks without first quitting their current task.

Each agent-based simulation lasted *T* = 10,000 time steps. All simulations and the subsequent analyses were conducted in R (79).

## Acknowledgments

We thank A. Gal for advice on analyses, and O. Feinerman and M. Liu for contributions to the tracking algorithms. Research reported in this publication was supported by grants from the Faculty Scholars Program of the Howard Hughes Medical Institute, the Pew Biomedical Scholars Program, and the National Institute of General Medical Sciences of the National Institutes of Health under Award Number R35GM127007 (D.J.C.K.); Swiss National Science Foundation Advanced Postdoc Mobility (P300P3-147900) and Ambizione (PZ00P3_168066) fellowships, and a Rockefeller University Women & Science fellowship (Y.U.); Army Research Office Grant W911NF-18-1-0325 (M.K.); National Science Foundation Graduate Research Fellowship no. DGE1656466 (C.K.T.); and a Kravis Fellowship (J.S.). The content is solely the responsibility of the authors and does not necessarily represent the official views of the National Institutes of Health, the Howard Hughes Medical Institute, or the Pew Biomedical Scholars Program. This is Clonal Raider Ant Project paper #13.

