## Supplementary Information for "Emergent behavioral organization in heterogeneous groups of a social insect"

##### This PDF file includes:

Supplementary text
Figs. S1 to S8
Tables S1 to S2
References for SI reference citations

### Supporting Information Text

#### 1. Image acquisition and ant detection

To acquire behavioral data, we used an automated scan-sampling approach, in which a picture of each colony was acquired every ca. 400 seconds throughout the experiment by a custom setup comprising 28 webcams (Logitech® B910 or C910) and controlled LED lighting. Each webcam acquired images (5 megapixels, RGB) of four colonies, and the position of colonies within the setup was randomized.

Size and color-probability thresholds were used to detect candidate image regions corresponding to ants carrying color tags. Candidate ants were oriented and identified on the basis of the relative position of cuticle and tag color probability maxima along the main axis of each region. Based on a previous assessment (1), the ant identification algorithm correctly identifies on average 77.1% of the ants that can be manually identified (i.e., 22.9% of ants are missed). Of all the automated assignments, 94.4% on average are correct (i.e., 5.6% assign the wrong ID).

#### 2. Analytical treatment of the FTM

In addition to simulating the fixed threshold model (FTM) described in Materials and Methods, we derive analytical predictions for the long-term behavior of the FTM. Comparing these predictions with simulation results furthers our understanding of what mechanisms give rise to the behavioral patterns observed in the experimental data. Below is a brief summary of the key equations of the model as well as the simplifying assumptions we make in order to gain analytical insight; we refer the reader to Materials and Methods for details of the model.

Recall that the model considers a colony of $n$ individuals performing $m$ tasks; we assume that there are two tasks ($m = 2$; see Materials and Methods). Individuals can be of one of two types, X or Y. To analytically study how individual behavior depends on the ratio of the types, we define $f$ and $1 - f$ to be the fractions of the colony consisting of individuals of type X and Y, respectively.

The model assumes that individual $i$'s internal threshold $\theta_{ij}$ is drawn from a normal distribution with mean $\mu_j$ and normalized standard deviation $\sigma_j$. For our analytical analysis, we make the simplifying assumption that $\sigma_j = 0$ for all tasks. In other words, we assume that the type- and task-specific thresholds are given by the constant parameters, $\mu_j^X$ and $\mu_j^Y$. With this assumption, the probabilities $P_{ij,t}^X$ and $P_{ij,t}^Y$ that inactive individuals $i$ of types X and Y begin to perform task $j$ at time $t$ are, respectively,

$$P_{j,t}^X\left(s_{j,t}\right) = \frac{s_{j,t}^\eta}{s_{j,t}^\eta + \left(\mu_j^X\right)^\eta}, \qquad P_{j,t}^Y\left(s_{j,t}\right) = \frac{s_{j,t}^\eta}{s_{j,t}^\eta + \left(\mu_j^Y\right)^\eta}. \qquad [1]$$

Because we assume that there are two tasks ($m = 2$), the numbers of X and Y individuals performing task $j$ at time $t + 1$ are governed by the following equations, in which $j'$ denotes the other task:

$$\begin{aligned} n_{j,t+1}^X - n_{j,t}^X &= \frac{1}{2}\left[P_{j,t}^X(s_{j,t}) + \left(1 - P_{j',t}^X\left(s_{j',t}\right)\right) P_{j,t}^X\left(s_{j,t}\right)\right]\left(fn - \left(n_{j,t}^X + n_{j',t}^X\right)\right) - \tau n_{j,t}^X \\ n_{j,t+1}^Y - n_{j,t}^Y &= \frac{1}{2}\left[P_{j,t}^Y(s_{j,t}) + (1 - P_{j',t}^Y(s_{j',t}))P_{j,t}^Y(s_{j,t})\right]\left((1-f)n - \left(n_{j,t}^Y + n_{j',t}^Y\right)\right) - \tau n_{j,t}^Y, \end{aligned} \qquad [2]$$

where $n_{j,t}^X$ and $n_{j,t}^Y$ are the numbers of X and Y individuals performing task $j$ at time $t$, respectively, and $\tau$ is the probability of quitting a task. The sums in the larger parentheses represent the pool of individuals who could possibly initiate task $j$—that is, the total number of inactive individuals. The sums in square brackets then capture the possible ways in which these inactive individuals can initiate task $j$: they can either encounter the stimulus for task $j$ immediately and begin performing that task, or they can first encounter the stimulus for the other task $j'$, not perform that task, subsequently encounter the stimulus for task $j$, and begin performing task $j$.

Finally, recall that the dynamics of the stimulus $s_{j,t}$ associated with task $j$ is governed by Eq. (3):

$$s_{j,t+1} - s_{j,t} = \delta_j - \frac{\alpha_j^X n_{j,t}^X + \alpha_j^Y n_{j,t}^Y}{n}, \qquad [3]$$

where $\delta_j$ is the task-specific demand rate, and $\alpha_j^X$ and $\alpha_j^Y$ are the task-specific performance efficiencies of X and Y individuals, respectively.

In the subsequent sections, we compute the long-term behavior of the system of six difference equations in Eq. (2) (for $n_1^X, n_1^Y, n_2^X, n_2^Y$) and Eq. (3) (for $s_1, s_2$) and compare the results to simulations.

##### A. Steady-state predictions.

***A.1. Theoretical maximum activity level.*** In the model, individuals have a latency period of one time step between when they quit a task and when they recommence working. This means that, on average, only a fraction of the colony can be working at any given time.

To find this maximal activity level, let $Z_t = (n_{1,t}^X + n_{1,t}^Y + n_{2,t}^X + n_{2,t}^Y)/n$ be the fraction of active individuals in a colony at time $t$. Note that $0 \leq Z_t \leq 1$. At time $t + 1$, on average, a fraction $\tau Z_t$ of the colony becomes inactive. Therefore, at time $t + 1$,

$$(\text{fraction active}) + (\text{fraction inactive}) = X_{t+1} + \tau Z_t \leq 1.$$

At steady state, the equality $X_{t+1} = X_t = Z^*$ is satisfied. By substitution, we obtain the theoretical maximum activity level:

$$Z^* \leq \frac{1}{1+\tau}. \qquad [4]$$

For example, for $\tau = 0.2$ used in the simulations (see Fig. 4), at most 83.33% of the individuals in a colony can be active at steady state. A similar condition has been noted by (2).

**A.2. Pure colonies.** Without loss of generality, we consider pure colonies that consist of type X individuals only: let $f = 1$ and $n^Y_{j,t} = 0$ for all $t$. By setting and Eq. (3) to zero, we obtain the fraction of X individuals performing task $j$ at steady state, given by

$$\frac{n_j^X}{n} = \frac{\delta_j}{\alpha_j^X}. \qquad [5]$$

Notably, the steady-state values of $n_j^X$ are independent of the mean threshold ($\mu_j^X$) or the quit probability ($\tau^X$). This agrees with our simulation results in which differences in $\mu$ (Fig. 1c) or $\tau$ (Fig. S2b) alone did not change the mean task performance levels in pure colonies.

According to the condition in Eq. (4), this steady state is biologically possible only if

$$(Z^* =)\ \frac{n_1^X}{n} + \frac{n_2^X}{n} = \frac{\delta_1}{\alpha_1^X} + \frac{\delta_2}{\alpha_2^X} \leq \frac{1}{1+\tau}.$$

If this condition is not met, then we would expect the stimuli to continue growing (i.e., the system would not reach a steady state).

Now, suppose that the demand rate and task performance efficiency are the same for both tasks ($\delta_1 = \delta_2 = \delta$, $\alpha_1^X = \alpha_2^X = \alpha^X$). Eq. (5) implies that the fractions of X individuals performing tasks 1 and 2 at steady state would be

$$\frac{n_1^X}{n} = \frac{n_2^X}{n} = \frac{\delta}{\alpha^X}.$$

Similarly, in pure colonies of type Y, if $\delta_1 = \delta_2 = \delta$ and $\alpha_1^Y = \alpha_2^Y = \alpha^Y$, then $n_1^Y/n = n_2^Y/n = \delta/\alpha^Y$ at steady state. Thus, in order for pure colonies of type X and type Y to have different average task performance levels (i.e., $\delta/\alpha^X \neq \delta/\alpha^Y$) under the assumptions above—i.e., that the tasks are equally demanding and that a given type of individual is equally efficient at both tasks—the two types must differ in task performance efficiency ($\alpha^X \neq \alpha^Y$) (see Results).

**A.3. Mixed colonies with 1:1 mixes.** We now consider mixed colonies consisting of X and Y individuals in equal proportions ($f = 0.5$). We assume that the mean thresholds and the quit probabilities are identical for both tasks and ant types ($\mu_1^X = \mu_2^X = \mu_1^Y = \mu_2^Y$ and $\tau^X = \tau^Y$)*. Setting Eq. (2) and Eq. (3) equal to zero, we find that the steady-state numbers of individuals performing task $j$ are given by

$$n_j^X = n_j^Y = n\left(\frac{\delta_j}{\alpha_j^X + \alpha_j^Y}\right).$$

This quantity can also be expressed as a fraction of each type of individuals:

$$\frac{n_j^X}{(n/2)} = \frac{n_j^Y}{(n/2)} = \frac{2\delta_j}{\alpha_j^X + \alpha_j^Y}. \qquad [6]$$

Applying condition Eq. (4), this steady-state is only biologically relevant when

$$(Z^* =)\ \sum_{j=1}^{2} \frac{n_j^X}{n} + \frac{n_j^Y}{n} = \sum_{j=1}^{2} \frac{2\delta_j}{\alpha_j^X + \alpha_j^Y} \leq \frac{1}{1+\tau}.$$

Again, if this condition is not met, then we would expect the stimuli to continue growing over time and for the individuals to be working at maximum capacity.

**A.4. Mixed colonies with non-1:1 mixes.** We now generalize to the case in which a fraction $f$ of individuals ($0 \leq f \leq 1$) in a mixed colony are of type X. In the simplified case where $\mu_1^X = \mu_2^X = \mu_1^Y = \mu_2^Y$ and $\tau^X = \tau^Y$, the steady-state fractions of individuals performing task $j$ are

$$n_j^X = \frac{fn\delta_j}{f\alpha_j^X + (1-f)\alpha_j^Y}, \quad n_j^Y = \frac{(1-f)n\delta_j}{f\alpha_j^X + (1-f)\alpha_j^Y}.$$

Since there are $fn$ individuals of type X and $(1-f)n$ individuals of type Y, these quantities can be expressed as fractions of individuals of type X and Y individuals performing task $j$:

$$\frac{n_j^X}{fn} = \frac{n_j^Y}{(1-f)n} = \frac{\delta_j}{f\alpha_j^X + (1-f)\alpha_j^Y} \left(= \frac{n_j^X + n_j^Y}{n}\right). \qquad [7]$$

The last equality highlights the fact that, at steady state, the fraction of individuals of each type performing task $j$ is identical to the fraction of the whole colony performing that task, i.e., both X and Y perform task $j$ at equal rates. As expected, the expressions Eq. (7) reduce to Eq. (6) when $f = 0.5$ (1:1 mixes) and to Eq. (5) when $f = 1$ (pure colonies with X individuals only). Again, we expect to see this equilibrium only when condition (4) is satisfied.

From Eq. (7), we expect the steady-state task $j$ performance frequency to depend non-linearly on the fraction of X individuals ($f$).

*The parameters $\mu$ and $\tau$ do not explicitly appear in Eq. (6) when we assume that the mean thresholds are identical for all individuals and both tasks. However, based on the Eq. (2), we expect the general form of steady state fractions of active individuals to be explicit functions of $\mu_j^X$ and $\mu_j^Y$ as well as $\tau^X$ and $\tau^Y$. While the steady states can be computed numerically for the case when these parameters differ between types or tasks, the analytical expressions are too complicated to write down.

***A.5. Mixed colonies with symmetric mean thresholds.*** So far we have assumed that the mean task thresholds $\mu_j^X$ and $\mu_j^Y$ are identical for both ant types and tasks ($\mu_1^X = \mu_2^X = \mu_1^Y = \mu_2^Y$). While the system of equations (Eq. (2) and Eq. (3)) can be solved numerically even when we introduce between-type differences in $\mu$, the steady-state expressions quickly become too difficult to write down. In the following special case, however, we can express the equilibrium values exactly. Assume that

1. colonies consist of type X and Y individuals in equal proportions ($f = 0.5$);
2. task efficiency is the same for both ant types and tasks ($\alpha_1^X = \alpha_2^X = \alpha_1^Y = \alpha_2^Y = \alpha$);
3. demand rate is the same for both tasks ($\delta_1 = \delta_2 = \delta$); and
4. mean task thresholds are symmetric, such that one type has a low threshold for one task and a high threshold for the other while this ordering is reversed in the other type: $\mu_1^X = \mu_2^Y = a$ and $\mu_2^X = \mu_1^Y = b$.

Importantly, the symmetry between the two tasks and between the two types imply that the stimulus levels for the tasks would be identical at stead state ($s_1 = s_2 = s^*$). Moreover, at steady state, the number of X individuals performing task 1 would be identical to the number of Y individuals performing task 2 ($n_1^X = n_2^Y$); similarly, we would expect that $n_1^Y = n_2^X$. Substituting these conditions into Eq. (3) and setting it equal to zero, we find that, at steady state,

$$n_1^X + n_1^Y = n_2^X + n_2^Y = n_1^X + n_2^X = n_1^Y + n_2^Y = n\left(\frac{\delta}{\alpha}\right).$$

By substituting this into Eq. (2) and following the symmetry argument above, we derive an expression for the steady-state stimulus level $s^*$:

$$s^*(= s_1 = s_2) = \left[\frac{1}{2}\left(-(a^\eta + b^\eta) \pm \sqrt{(a^\eta + b^\eta)^2 + (a^\eta b^\eta) \cdot \frac{8\delta\tau}{\alpha - 2\delta(1+\tau)}}\right)\right]^{\frac{1}{\eta}}.$$

The corresponding steady-state fractions of X and Y individuals performing tasks 1 and 2 are

$$\frac{n_1^X}{(n/2)} = \frac{n_2^Y}{(n/2)} = \frac{1}{\tau}\left(\frac{(s^*)^\eta}{(s^*)^\eta + a^\eta}\right)\left[2 - \frac{(s^*)^\eta}{(s^*)^\eta + b^\eta}\right]\left(\frac{1}{2} - \frac{\delta}{\alpha}\right),$$
$$\frac{n_2^X}{(n/2)} = \frac{n_1^Y}{(n/2)} = \frac{1}{\tau}\left(\frac{(s^*)^\eta}{(s^*)^\eta + b^\eta}\right)\left[2 - \frac{(s^*)^\eta}{(s^*)^\eta + a^\eta}\right]\left(\frac{1}{2} - \frac{\delta}{\alpha}\right). \quad [8]$$

When $a = b$ (i.e., when all $\mu$'s are the same), these expressions reduce to the steady-state values predicted in Eq. (6).

#### B. Comparing analytical predictions with simulation results.

***B.1. Steady-state prediction 1: differences in task efficiency and demand rate.*** Overall, our predictions perform well against the simulations. Figure S6 gives illustrative comparisons for two cases in which the two ant types differ in task efficiency only. The cases in turn differ in demand rate only. When colonies can keep up with the demand—i.e., the demand rate is sufficiently low relative to the task efficiencies of both types such that the system satisfies Eq. (4)—there is strong agreement between our analytical predictions (Eq. (6)) and simulation results for the steady-state task performance (Fig. S6a). In contrast, when the demand rate is too high such that the colony cannot keep up with the demand, colonies violate the necessary condition for reaching a steady state (Eq. (4)); indeed, we observe that the stimuli run away (i.e., keep increasing over time) in the simulations, an indication that the system is not in equilibrium. As expected, our analytical predictions in Eq. (6) do not match the simulation results in this case (Fig. S6b).

***B.2. Steady-state prediction 2: symmetric mean thresholds.*** Figure S7 compares our analytical predictions (Eq. (8)) and simulations for two cases with symmetric thresholds: that is, the mean threshold $\mu$ differs symmetrically between ant types ($\mu_1^X = \mu_2^Y$ and $\mu_2^X = \mu_1^Y$) and all other parameters are the identical for both types and tasks. Both cases (Fig. S7a-b) show strong agreement between the simulated and analytically-predicted results for steady-state task performance.

##### C. Downward vs. upward contagion.

Both our experiments (Fig. 2a-b) and theoretical analyses (Fig. 4a-b) demonstrated patterns of asymmetric behavioral contagion between the types, in which individuals of different types were behaviorally more similar to each other when mixed. Here we combine our analytical predictions for pure and mixed colonies to investigate conditions under which such contagion patterns arise. Consider two pure colonies consisting of X and Y individuals, respectively, and a third, mixed colony consisting of a 1:1-ratio of X and Y individuals. Let us assume that each colony reaches a steady state (i.e., Eq. (4) is satisfied for each colony). We show analytically that, under these conditions, if the ant types only differ in task efficiency ($\alpha_j^X, \alpha_j^Y$), then the system can exhibit a downward contagion but not an upward contagion.

We can directly apply the steady-state fractions of active individuals computed in Eq. (5) and Eq. (6) because the mean threshold ($\mu$) and the quit probability ($\tau$) are assumed to be identical across ant types. The behavioral contagion is downward (Fig. S8a) if

$$\frac{1}{2}\left(\frac{\delta_j}{\alpha_j^X} + \frac{\delta_j}{\alpha_j^Y}\right) > \frac{2\delta_j}{\alpha_j^X + \alpha_j^Y} \quad [9]$$

and upward (Fig. S8b) if the inequality is reversed (see also Fig. 4a-b).

By manipulating the inequality Eq. (9), we see that the left-hand side is always at least as large as the right-hand side:

$$\frac{1}{2}\left(\frac{\delta_j}{\alpha_j^X} + \frac{\delta_j}{\alpha_j^Y}\right) - \frac{2\delta_j}{\alpha_j^X + \alpha_j^Y} = \frac{\delta_j}{2}\left(\frac{(\alpha_j^X - \alpha_j^Y)^2}{\alpha_j^X \alpha_j^Y (\alpha_j^X + \alpha_j^Y)}\right) \geq 0$$

The equality holds if and only if $\alpha_j^X = \alpha_j^Y$, in which case the types are indistinguishable with respect to task $j$. If $\alpha_j^X \neq \alpha_j^Y$, then only downward contagion is possible under our assumptions. Note that this result is agnostic to between-task differences in task efficiency or task demand; in other words, it holds even when $\alpha_1^X \neq \alpha_2^X$, $\alpha_1^Y \neq \alpha_2^Y$, and $\delta_1 \neq \delta_2$.

**Table S1. List of experimental treatments.** Text in bold denotes the variable of interest for each experiment. All mixed colonies contained a 1:1 ratio of each ant type.

| Experiment | Worker genotype | Brood genotype | Age (cycles) | Subcaste | Colony size | # replicates | # colonies |
|---|---|---|---|---|---|---|---|
| Genetic composition 1 | **A, B, mixed** | A | 1 | Regular workers | 16 | 8 | 24 |
| Genetic composition 2 | **A, B, mixed** | B | 1 | Regular workers | 16 | 8 | 24 |
| Demographic composition | B | B | **1, 3, mixed** | Regular workers | 16 | 8 | 24 |
| Morphological composition | B | B | 1 | **Regular workers, intercastes, mixed** | 8 | 16 | 48 |

**Table S2. Parameter settings for fixed threshold model (FTM).**

| *Parameters* | *Description* | *Values in simulations* |
|---|---|---|
| $T$ | Simulation length in time steps | 10,000 |
| $n$ | Number of individuals | 8, 16 |
| $m$ | Number of tasks | 2 |
| $\delta_j = \delta$ | Brood-specific rate of stimulus increase (i.e., demand rate); taken to be the same for all tasks | 0.6, 1.3 |
| $\alpha_j^X = \alpha^X$ $\alpha_j^Y = \alpha^Y$ | Type-specific performance efficiency of active individuals for task $j$; taken to be the same for all tasks | 1 – 6 |
| $\mu_j^X = \mu^X$ $\mu_j^Y = \mu^Y$ | Mean of the type-specific threshold distribution for task $j$; taken to be the same for all tasks | 6.5 – 15.5 |
| $\sigma_j^X = \sigma^X$ $\sigma_j^Y = \sigma^Y$ | Variance of the type-specific threshold distribution for task $j$ as a fraction of the corresponding mean; taken to be the same for all tasks | 0.1, 0.5 |
| $\eta$ | Threshold stochasticity | 7 |
| $\tau_j^X = \tau^X$ $\tau_j^Y = \tau^Y$ | Type-specific probability of quitting task $j$ once active (inverse of average task performance duration); taken to be the same for all tasks | 0.2, 0.6 |

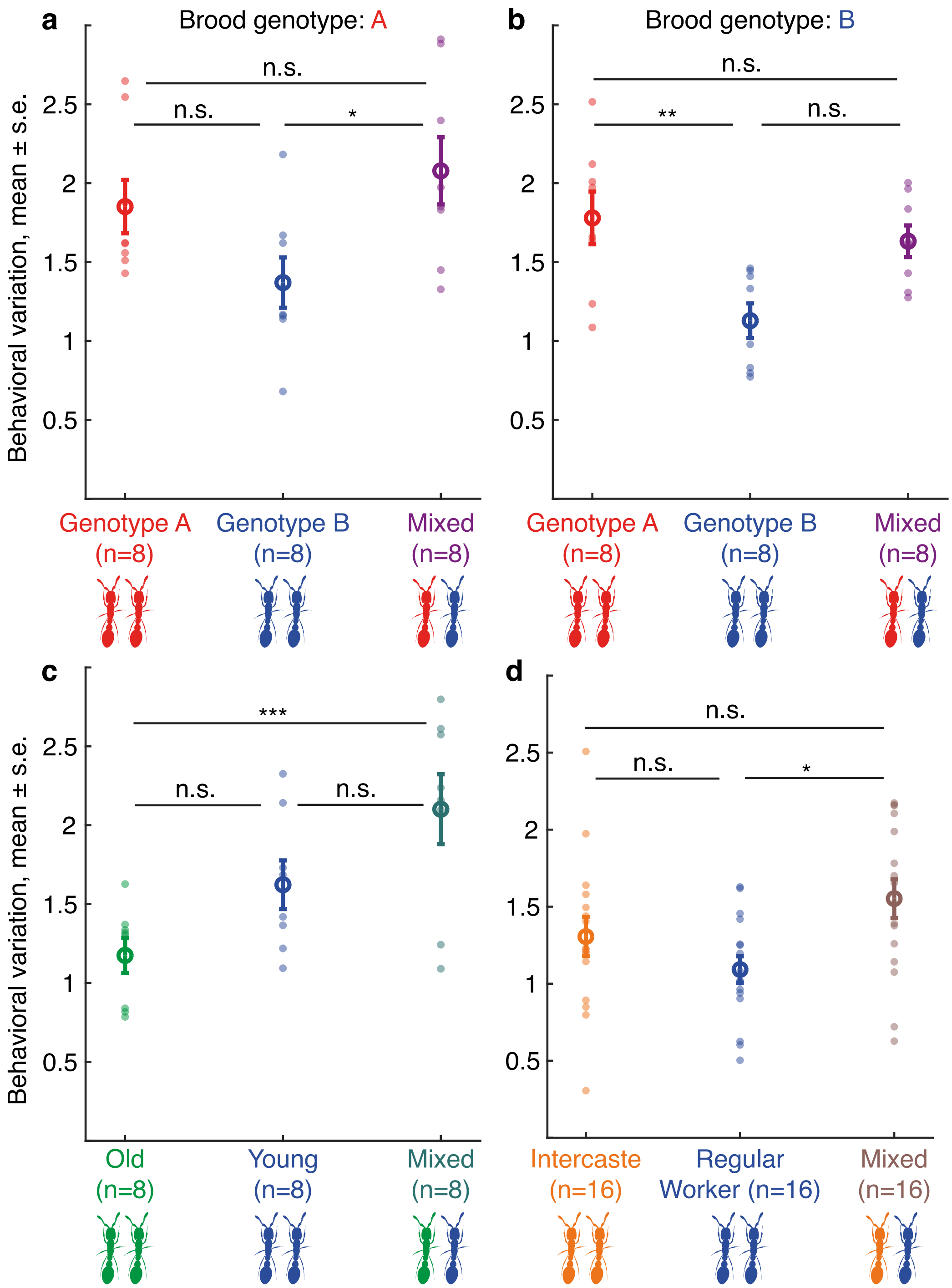


**Fig. S1. Behavioral variation (standard deviation in r.m.s.d. across colony members) as a function of colony composition.** Small full circles represent individual colonies; large open circles represent the average values across replicate colonies. Sample sizes indicate the number of replicate colonies. Identical colors across panels indicate ants of the same genotype, age, and morphological types. **a**: Behavioral variation as a function of colony genetic composition in colonies with A brood. Colony size 16 ($B_{hom}$ vs. Mixed: $z = -2.85$, $p = 0.013$; $A_{hom}$ vs. Mixed: $z = 0.81$, $p = 0.42$). **b**: Behavioral variation as a function of colony genetic composition in colonies with B brood. Colony size 16 ($B_{hom}$ vs. Mixed: $z = -2.15$, $p = 0.09$; $A_{hom}$ vs. Mixed: $z = 1.13$, $p = 0.52$). **c**: Behavioral variation as a function of colony demographic composition. Colony size 16 ($Young_{hom}$ vs. Mixed: $z = 1.81$, $p = 0.09$; $Old_{hom}$ vs. Mixed: $z = 3.84$, $p = 3.77 \cdot 10^{-04}$). **d**: Behavioral variation as a function of colony morphological composition. Colony size 8 (Regular $Worker_{hom}$ vs. Mixed: $z = -2.68$, $p = 0.02$, $Intercaste_{hom}$ vs. Mixed: $z = 1.47$, $p = 0.28$). n.s.: non-significant, *: $p < 0.05$, **: $p < 0.01$, ***: $p < 0.001$.

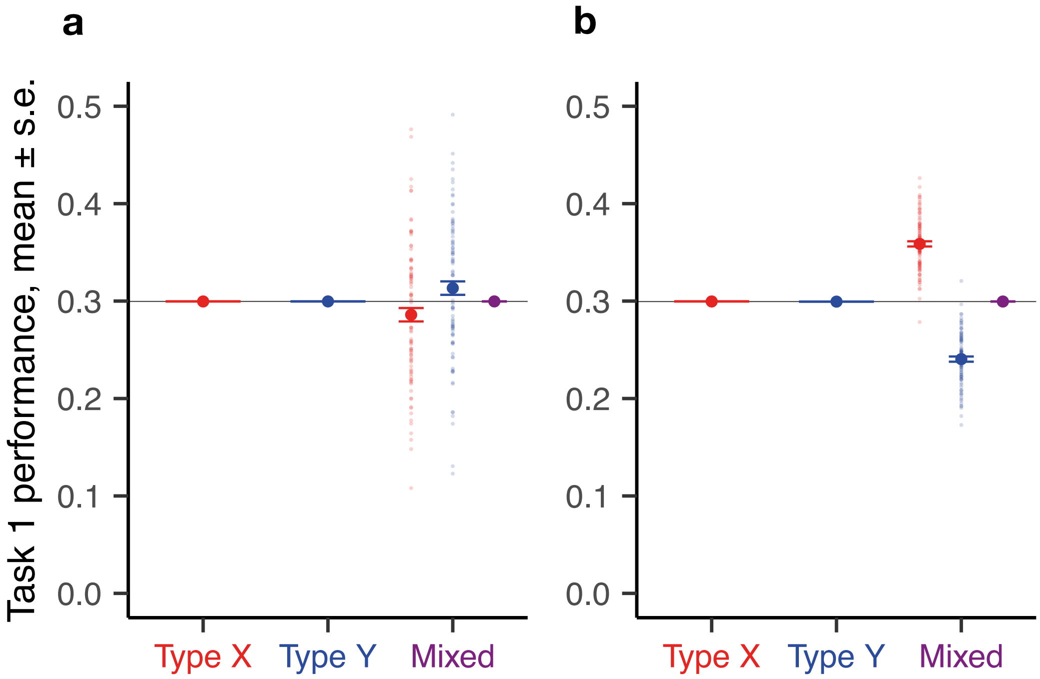


**Fig. S2. Theoretical predictions of the FTM with differences in threshold variance or task performance duration only.** Task performance frequency as a function of colony composition. One hundred replicates were simulated for each colony composition. Each opaque circle represents an individual replicate colony; each solid circle represents the average value across all replicates for its corresponding colony (or sub-colony) composition. **a**: Y individuals have a higher threshold variance than X individuals for both tasks ($\sigma^X$ = 0.1, $\sigma^Y$ = 0.5). **b**: Y individuals have a higher quitting rate (i.e., a shorter average task performance duration) than X individuals for both tasks ($\tau^X$ = 0.2, $\tau^Y$ = 0.6). All other parameters are identical for both types: $\mu$ = 10, $\delta$ = 0.6, $\alpha$ = 2, $\eta$ = 7.

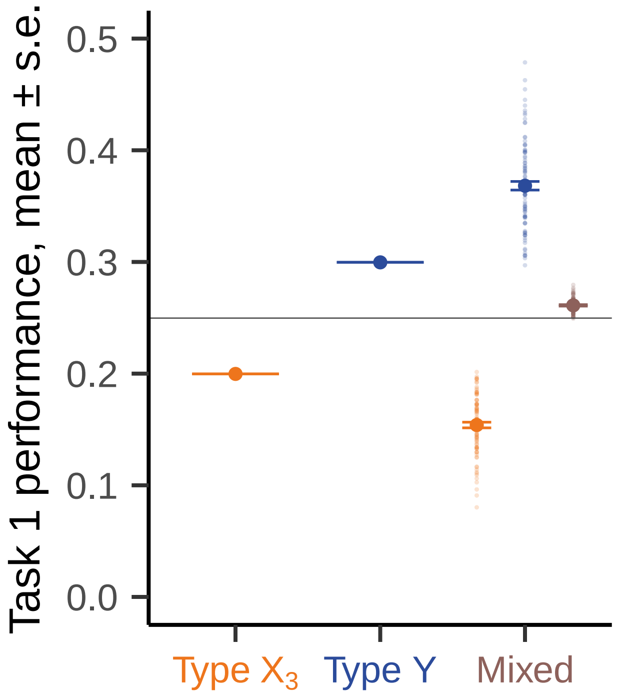


**Fig. S3. Theoretical predictions of the FTM on task performance for colony size 8.** Task performance frequency as a function of colony composition. One hundred replicates were simulated for each colony composition. Each replicate consisted of 8 individuals. Parameters are identical to those in Fig. 4d in the main text. The behavioral amplification pattern observed in Fig. 4d is recovered.

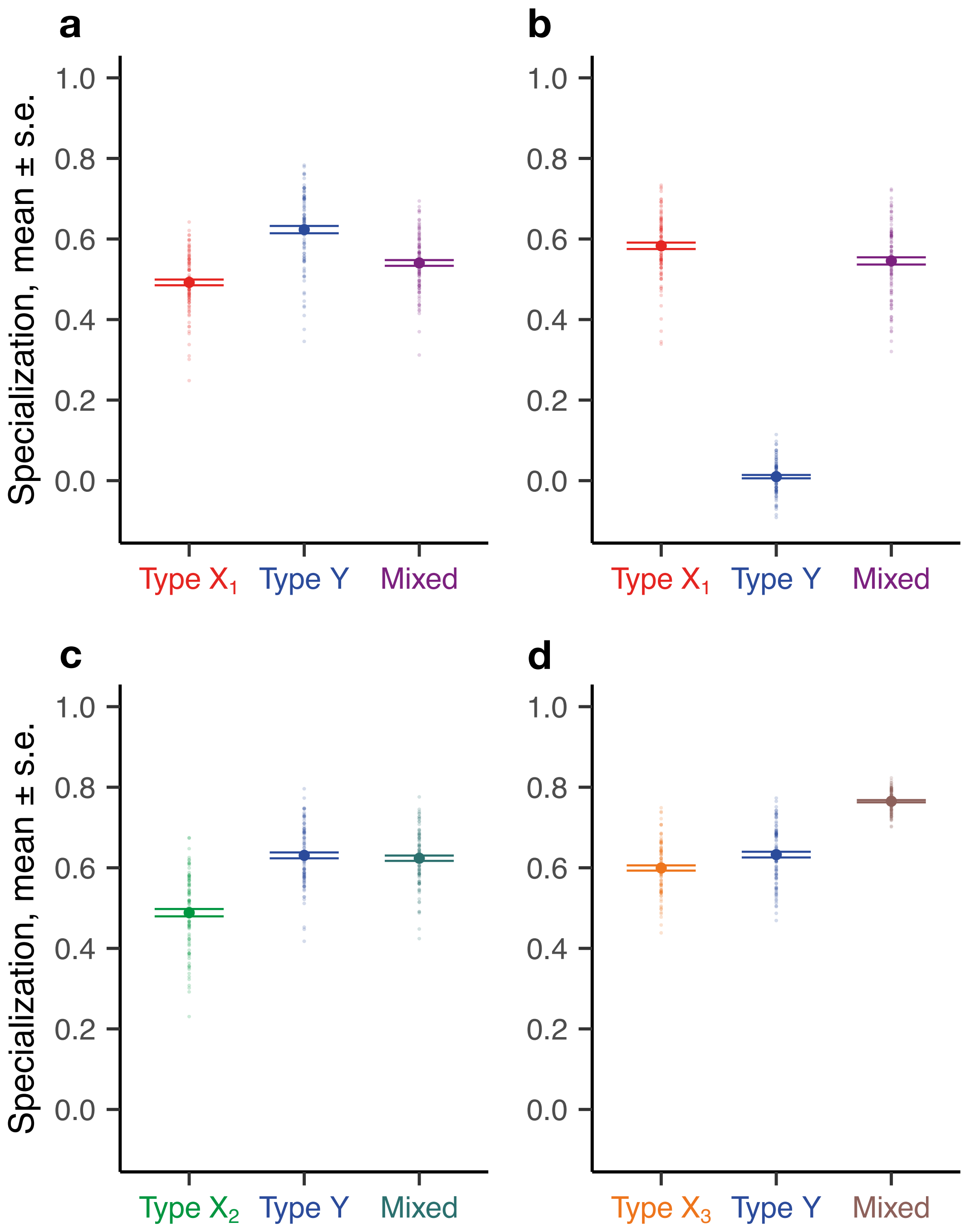


**Fig. S4. Theoretical predictions of the FTM on behavioral specialization.** Specialization was quantified using Spearman rank correlation on consecutive windows of 200 time steps. One hundred replicates were simulated for each colony composition (colony size 16). Each opaque circle represents an individual replicate colony; each solid circle represents the average value (mean ± s.e.m.) across all replicates for its corresponding colony composition. Identical colors and labels ($X_1$, $X_2$, $X_3$, Y) in this figure and Fig. 4 indicate individuals of the same type. Parameters are as in Fig. 4a-d.

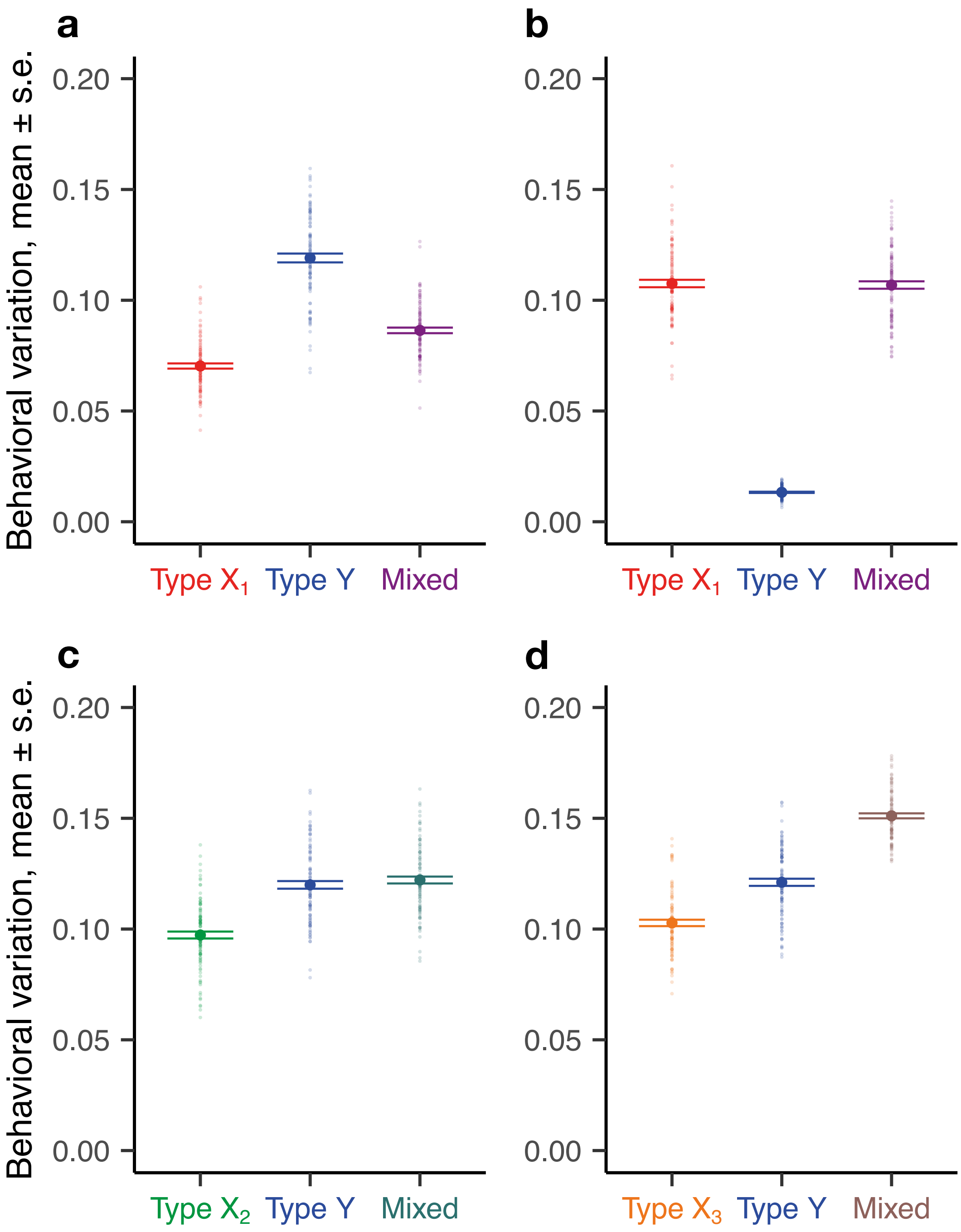


**Fig. S5. Theoretical predictions of the FTM on behavioral variation.** Behavioral variation was quantified as the standard deviation of task performance frequency across individuals in a colony. One hundred replicates were simulated for each colony composition (colony size 16). Each opaque circle represents an individual replicate colony; each solid circle represents the average value (mean ± s.e.m.) across all replicates for its corresponding colony composition. Identical colors and labels ($X_1$, $X_2$, $X_3$, Y) in this figure and Fig. 4 indicate individuals of the same type. Parameters are as in Fig. 4a-d.

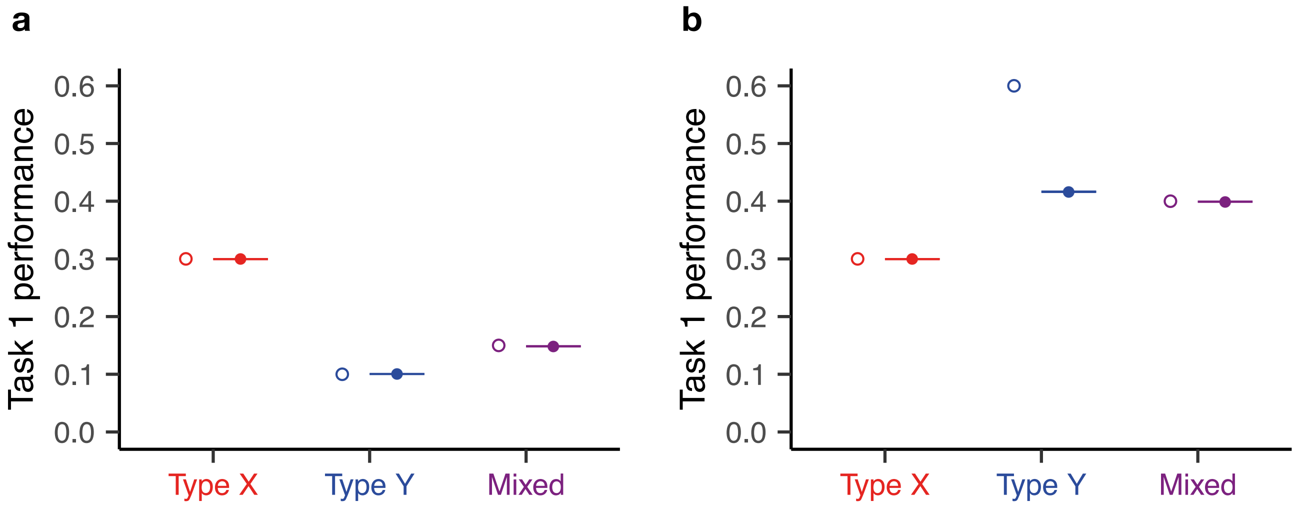


**Fig. S6. Comparison of simulation results and analytical predictions of steady states with differences in task efficiency.** Task performance frequency as a function of colony composition. One hundred replicates (colony size 16) were simulated for each colony composition and for each parameter combination. Each solid circle represents the average value (±s.e.m.) across all replicates for the corresponding colony composition; each empty circle represents the analytical prediction for that composition. Identical colors across panels indicate individuals of the same type. **a**: Both X and Y individuals can keep up with the demand ($\alpha_j^X$ = 2, $\alpha_j^Y$ = 6). **b**: X individuals can keep up with the demand ($\alpha_j^X$ = 2) but Y individuals cannot ($\alpha_j^Y$ = 1). Parameters: $\mu$ = 10, $\sigma$ = 0, $\eta$ = 7, $\delta$ = 0.6.

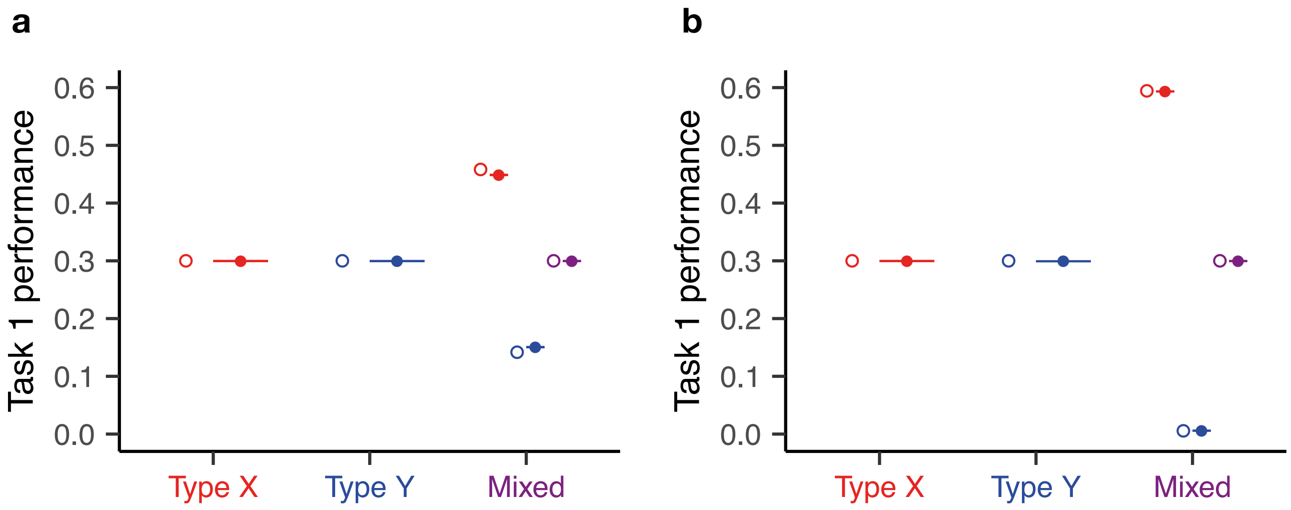


**Fig. S7. Comparison of simulation results and analytical predictions of steady states with symmetric mean thresholds.** Task performance frequency as a function of colony composition. One hundred replicates (colony size 16) were simulated for each colony composition and for each parameter combination. Each solid circle represents average value (±s.e.m.) across all replicates for the corresponding colony (or sub-colony) composition; each empty circle represents the analytical prediction for that colony (or sub-colony) composition. For both **a** and **b**, $\mu_1^X = \mu_2^Y = 10$; for **a**, $\mu_1^Y = \mu_2^X = 12$; for **b**, $\mu_1^Y = \mu_2^X = 12$. Parameters: $\sigma = 0$, $\eta = 7$, $\delta = 0.6$, $\alpha = 2$.

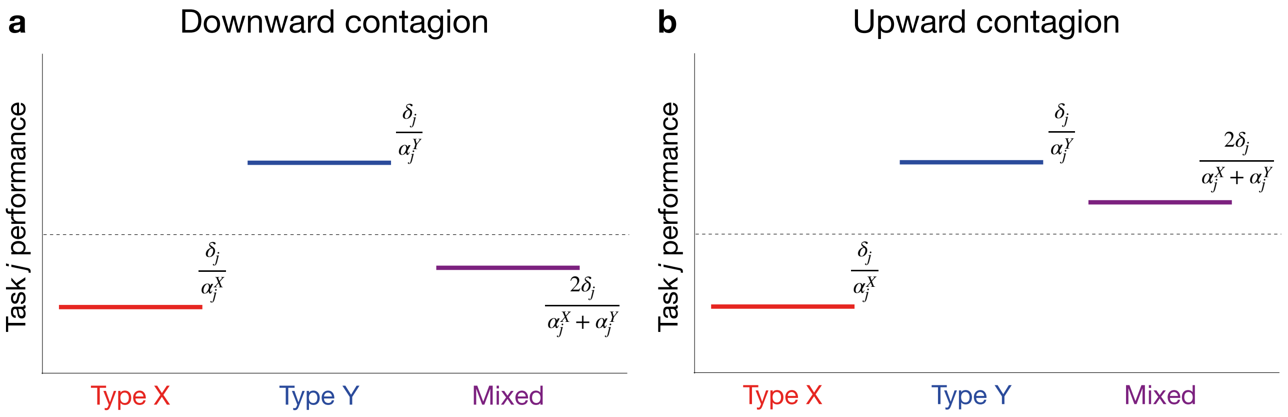


**Fig. S8. Schematic representation of the two asymmetric contagion patterns of our interest.** Downward (**a**) and upward (**b**) contagion illustrated in terms of task performance frequency as a function of colony composition. The mean thresholds and the quit probabilities are assumed to be identical for both ant types and both tasks ($\mu_1^X = \mu_2^X = \mu_1^Y = \mu_2^Y$ and $\tau^X = \tau^Y$). Without loss of generality, the figure also assumes that $\alpha_j^X > \alpha_j^Y$.
